# Molecular and Epigenetic Pathways Underlying Epithelial Damage and Repair in Necrotizing Enterocolitis via Multi-omics Approach

**DOI:** 10.1101/2025.04.17.647851

**Authors:** Yi Xiong, Andrea Zito, Haoyan Liang, George Biouss, Jielin Yang, Felicia Balsamo, Mina Yeganeh, Carol Lee, Dorothy Lee, Chen-Yi Wang, Nareh Tahmasian, Jinling Huang, Adam Minich, Mehrsa Feizi, Shiwen Wang, Yina Tian, Paolo De Coppi, Brian T. Kalish, Paul Delgado Olguin, Haitao Zhu, Bo Li, Agostino Pierro

**Author notes:** Equal contribution of first authors to this work (Yi Xiong, Andrea Zito, Haoyan Liang). Equal contribution corresponding authors to this work (Haitao Zhu, Bo Li, Agostino Pierro). **Correspondence: Haitao Zhu,** Department of General Surgery, Shanghai Children’s Hospital, School of Medicine, Shanghai Jiao Tong University, 355 Luding Road, Shanghai, 200062, China. Phone +86 21, 52976268.; **Bo Li,** Translational Medicine Program, The Hospital for Sick Children, 686 Bay Street, Toronto, ON M5G 1X8, Canada.; **Agostino Pierro,** Professor of Surgery, University of Toronto, Division of General and Thoracic Surgery, The Hospital for Sick Children, 1526-555 University Ave, Toronto, ON M5G 1X8, Canada.

## Abstract

**Introduction:** Neonatal necrotizing enterocolitis (NEC) is a severe gastrointestinal disorder with high mortality, characterized by epithelial cell injury and compromised epithelial repair. The mechanisms underlying defective epithelial regeneration remain poorly understood despite advances in single-cell omics. Addressing these challenges is essential for elucidating the pathogenesis of NEC and identifying therapeutic targets to restore epithelial regeneration and replace the damaged epithelial layer.

**Methods:** Multi-omics approaches were employed to investigate molecular and spatial changes in experimental NEC at epigenetic and transcriptomic levels. These included bulk RNA sequencing, single-nucleus RNA sequencing (snRNA-seq), single-nucleus assay for transposase-accessible chromatin sequencing (snATAC-seq), and multiplexed error-robust fluorescence in situ hybridization (MERFISH) for spatial transcriptomics. Complementary in vitro experiments and in vivo mouse models were utilized to evaluate NEC phenotypes, intestinal tissue morphology, and organoid formation.

**Results:** Changes in cell type composition, transcriptional network remodeling, and chromatin accessibility were observed in the small intestine of neonatal mice with NEC. Chromatin accessibility significantly changed in epithelial cells, highlighting their pivotal roles in NEC. A marked reduction in intestinal stem cells (ISCs) and transit-amplifying cells, along with an increased proportion of enteroendocrine cells, indicates disrupted epithelial regeneration and functional differentiation. These changes correlated with disrupted WNT signaling and stem cell maintenance genes (e.g., Lgr5, Smoc2, Axin2) and activation of inflammatory and hypoxia-related pathways (e.g., Il6, Tnfα). The epigenetic regulator Ezh2 was identified as a critical factor in maintaining LGR5+ ISCs and epithelial homeostasis. Knockdown of Ezh2 reduced stemness and proliferation-related gene expression and exacerbated inflammation. Reactivation of WNT signaling restored Ezh2 and Lgr5 expression, improving intestinal regeneration.

**Conclusion:** This study reveals dynamic transcriptomic, epigenetic, and spatial changes in NEC and highlights Ezh2 as a key regulator of LGR5+ intestinal stem cell function and epithelial regeneration. These findings provide insights into NEC pathogenesis and a basis for therapies targeting Ezh2 and WNT signaling to restore intestinal integrity.

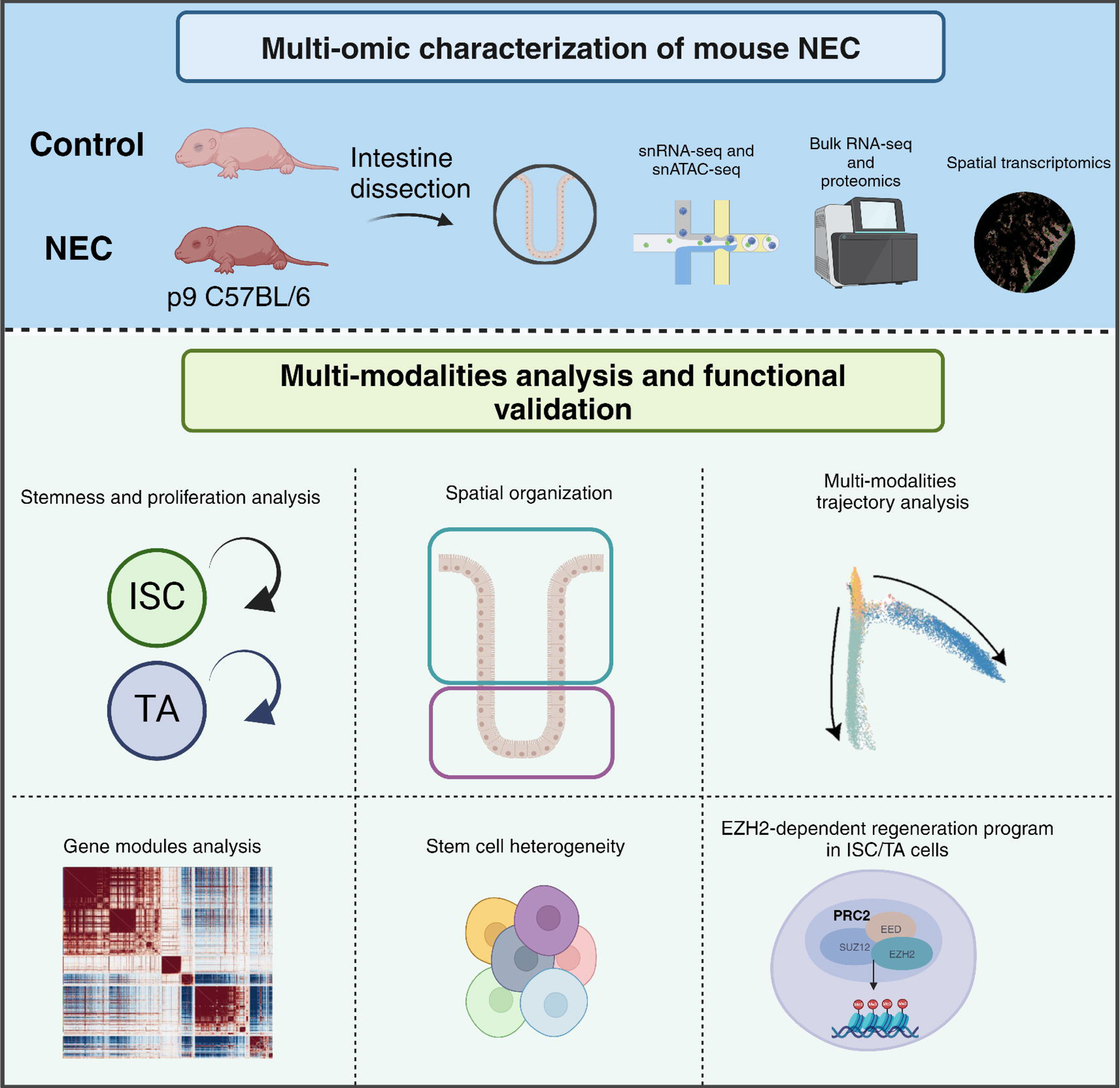

## Introduction

Necrotizing enterocolitis (NEC) is the most common cause of gastrointestinal emergency and death in premature infants. The disease is characterized by inflammation and necrosis of the intestine, most frequently in the terminal ileum. Complications range in severity from moderate intestinal inflammation to intestinal perforation, widespread bowel necrosis, multiple organ failure, and death [1]. NEC incidence ranges from 0.5 to 5 cases per 1000 live births [2], with significantly higher rates observed among very low birth weight and low-gestational age infants [3].

The intestinal epithelium plays a critical role in NEC pathogenesis due to its central function in maintaining gut integrity and barrier protection. This continuous layer of specialized cells, including absorptive enterocytes and secretory cells, not only facilitates digestion but also forms the first line of defense against luminal pathogens and toxins [4]. The compromised epithelium allows the translocation of bacteria and other harmful agents into the intestinal wall, triggering a cascade of inflammatory responses that lead to further tissue damage and necrosis [5]. However, the precise cellular mechanisms and specific epithelial cell types involved in NEC remain poorly understood, underscoring the need for advanced techniques to dissect these processes at a cellular and molecular level.

To maintain the integrity and optimal functioning of the intestinal epithelium, these specialized cells undergo constant turnover in a tightly controlled homeostatic process, replenishing the epithelium every 3-5 days [6]. This rapid renewal is driven by resident intestinal stem cells (ISCs) located in the intestinal crypts of Lieberkühn, which express Leucine-rich repeat-containing G-protein coupled receptor 5 (LGR5) [7]. LGR5+ ISCs undergo controlled division and give rise to highly proliferative transit-amplifying (TA) cells which migrate up the intestinal villus and differentiate into absorptive enterocytes or secretory cells [8, 9]. These cells replace less viable mature cells at the apex of intestinal villi that regularly undergo apoptosis. Evidence suggests that dysregulation of LGR5+ ISCs in NEC impairs the regeneration of intestinal epithelial tissue, correlating with severe gut damage and inflammation during NEC development [10–12]. However, the mechanism of LGR5+ ISC loss and whether ISC dysregulation is the cause or consequence during NEC remains uncharacterized.

Epigenetic transcriptional repression is critical for coordinating stem cell proliferation and differentiation to maintain tissue homeostasis. In humans, as in other multicellular organisms, polycomb group (PcG) proteins mediate transcriptional repression during development through two highly conserved multiprotein complexes: PRC1 and PRC2 [13]. PRC2 comprises four highly-conserved subunits, including EZH2 or its less-abundant isoform, EZH1, which serve as catalytic subunits possessing methyltransferase activity [14]. This activity catalyzes histone 3 lysine 27 (H3K27) mono-, di-, or trimethylation (H3K27me3), favoring a localized chromatin condensation that restricts access of transcriptional machinery to target genes [15]. PRC1, on the other hand, recognizes these H3K27me3 marks and reinforces transcriptional repression by ubiquitinating histone H2A, further compacting chromatin and ensuring stable silencing of target genes [16].

PRC2 targets genes involved in stem cell differentiation and proliferation, underscoring its pivotal role in these processes [13]. In adult mice, PRC2 maintains the ISC niche, promoting the proliferation of TA cells, and driving epithelial regeneration following intestinal damage [13]. Additionally, PRC2 restricts the differentiation of ISCs into secretory lineages, highlighting its role in maintaining ISC identity [13]. Notably, the PRC2 subunit, EZH2, has been shown to orchestrate homeostasis of various tissues through resident stem cell regulation, and its overexpression has been linked to uncontrolled cell growth in cancers, [17–20]. PRC2-EZH2-mediated transcriptional repression in ISCs during normal neonatal intestinal development, and in the context of NEC, remains unexplored.

This study aims to elucidate the epigenetic and transcriptomic mechanisms, particularly Ezh2-mediated pathways, that underlie intestinal stem cell dysregulation and impaired epithelial regeneration in necrotizing enterocolitis. Using a multi-omics approach, we identified significant reductions in epithelial cell proliferation and stemness markers—especially in enterocytes, intestinal stem cells (ISCs), and transit-amplifying (TA) cells—linked to dysregulated chromatin accessibility. Ezh2-mediated epigenetic mechanisms were implicated in regulatory networks affecting ISC maintenance and epithelial repair in NEC-related intestinal injury. These findings offer potential therapeutic targets for restoring intestinal homeostasis in NEC and contribute to our broader understanding of epigenetic regulation in gut health, with implications for other inflammatory and degenerative gastrointestinal diseases.

## Results

### Multi-omic atlas of the neonatal mouse intestine during NEC intestinal injury

To gain a comprehensive understanding of the molecular changes associated with NEC, we performed single-nucleus RNA sequencing (snRNA-seq) and single-nucleus assay for transposase-accessible chromatin sequencing (snATAC-seq) on the small intestine of an established neonatal mouse model of NEC at postnatal day P9 [12] (Fig. 1A).

**Figure 1.**
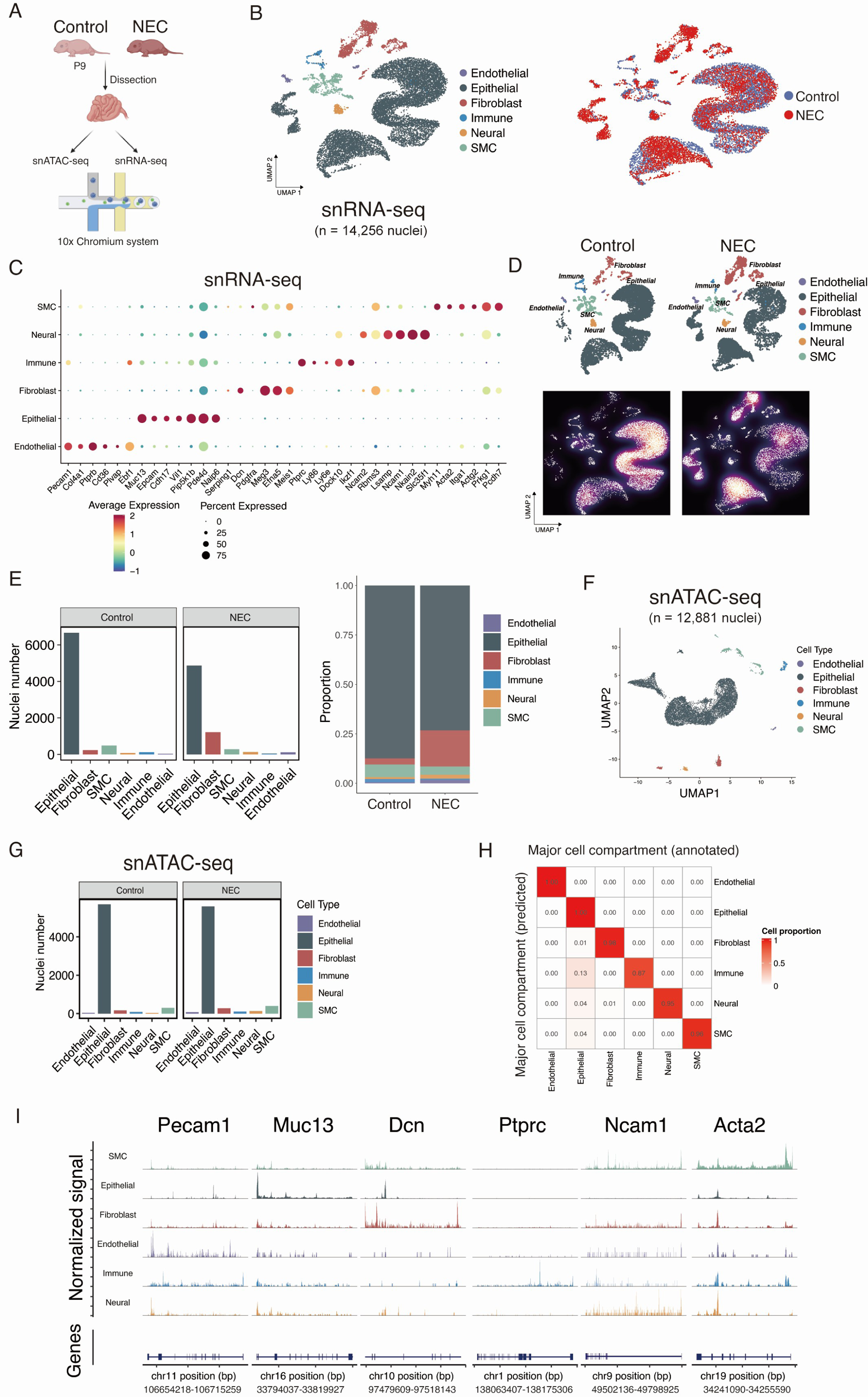
Multi-omic profiling of NEC intestine at the single-cell resolution. (A) Schematic diagram depicting the study design. (B) UMAP view of major cell types and sample group in snRNA-seq data. (C) Expression levels and frequencies of selected markers across major cell types. (D) UMAP view of major cell types separated by sample group. Galaxy plots depicting the cell density in UMAP for cells from the control (left) and NEC (right) groups. Cooler colors indicate low density, and warmer colors indicate high density. (E) Barplot depicting the cell number or proportion of major cell types in the control and NEC groups in snRNA-seq data. (F) UMAP view of major cell types in snATAC-seq data. (G) Barplot depicting the cell number of major cell types in the control and NEC groups in snATAC-seq data. (H) Heatmap depicting the correlation between manually annotated cell types and cell types predicted from Seurat workflow. (I) Normalized chromatin accessibility profiles for each cell type at canonical marker genes.

After rigorous quality filtering of the snRNA-seq data, we retained a total of 14,256 cells, comprising 7,620 cells from the control group and 6,636 cells from the NEC group (Fig. 1B). These cells passed quality control (QC) with an average of 696 genes per cell and 1,207 unique molecular identifiers (UMIs) per cell (Fig. S1A). To exclude technical batch effects, we corrected the datasets from all samples and tissues using Harmony [21]. Unsupervised clustering partitioned all cells into 25 clusters. Based on differential expression analysis and the expression of selected marker genes from the literature, we manually annotated the major cell types: epithelial (*Muc13*, *Epcam*, *Cdh17*), fibroblast (*Dcn*, *Meg3*, *Efna5*), neural (*Ncam1*, *Ncam2*, *Lsamp*), immune (*Ptprc*, *Ly86*, *Ly6e*), endothelial (*Pecam1*, *Col4a1*, *Ptprb*), and smooth muscle cells (SMCs) (*Myh11*, *Acta2*, *Itga1*) (Fig. 1C). The uniform manifold approximation and projection (UMAP) analysis revealed an altered proportion of major cell types between the control and NEC groups (Fig. 1D-E). Tissue distribution analyses showed that the immune and SMC compartment was decreased in NEC while the neural, endothelial, and fibroblast compartment was increased (P < 0.05, monte-carlo/permutation test) (Fig. S1C).

To understand the epigenetic changes in the ileum during NEC, we generated single-cell chromatin accessibility profiles. After filtering out low-quality nuclei, we obtained a total of 12,881 nuclei from all samples for snATAC-seq, with a median read depth of 13,943 fragments per nucleus and a median transcription start site (TSS) ratio of 4.77 (Fig. S1B). Among the 12,881 nuclei, 6,312 were from the control group and 6,569 were from the NEC group. After batch correction and unbiased clustering using Latent Semantic Analysis on term frequency-inverse document frequency (LSA-log (TF-IDF)), the nuclei formed 19 clusters (Fig. S1E). One cluster, comprising 574 nuclei, was due to low sequencing depth. For each gene, we computed gene activity scores by aggregating peaks per cell in the gene body and promoter region. We manually annotated the remaining 18 clusters into major cell types based on differential gene activity scores (Fig. 1F-H). To confirm major cell-type annotations in the snATAC-seq cell populations, we used a canonical correlation analysis (CCA)-based label-transfer algorithm from Seurat. This algorithm predicts cell types in snATAC-seq data using cell-type annotations from snRNA-seq cells as reference. We achieved a maximum prediction score of ≥0.8 in 98% of cells, demonstrating high correspondence between the two data modalities. The heatmap also revealed high correspondence between manually annotated cell types and predicted cell types (Fig. 1G). Normalized chromatin accessibility profiles for each cell type at canonical marker genes were also revealed (Fig. 1I).

To further investigate the differences in chromatin accessibility among all cell types, we performed MACS2 [22] to identify a total of 208,341 peaks representing potential *cis*-regulatory elements in our snATAC-seq data (Fig. S1F). We defined differentially accessible chromatin regions (DARs) as peaks exhibiting substantial differences among cell types (log2(fold change [FC]) > 0.25 and adjusted P < 0.05). We identified a total of 46,846 DARs, with approximately 10.5% (0.105 ± 0.027) of DARs closely associated with differentially expressed genes (DEGs) in their corresponding cell types. Notably, epithelial and smooth muscle cells (SMCs) exhibited the most DARs, while immune and endothelial compartments exhibited the least in NEC (Fig. S2A). The majority of DARs were located in the promoter, intronic, and distal regions of the genome, though the distribution of DARs varied across cell types (Fig. S2B-C). These findings imply that epithelial and smooth muscle cells undergo the most pronounced changes in chromatin accessibility, likely reflecting altered regulatory programs driving NEC pathology. In contrast, the immune and endothelial cells may play a more reactive or secondary role, responding to primary regulatory changes occurring in epithelial and smooth muscle cells.

To identify potential regulators across six major cell types, we performed chromVAR [23] and transcription factor footprinting analysis. This approach revealed enriched transcription factor binding activity, including FOXL2 in smooth muscle cells (SMC) [24], SMAD2/SMAD3 in epithelial cells [25], WT1 in fibroblasts, SOX17 in endothelial cells, RUNX1 in immune cells, and SOX3 in the neural cell compartment (Fig. S2D). These findings highlight a diverse footprint enriched across the dataset, reflecting the distinct transcriptional regulatory landscapes of the major cell types.

Altogether, our transcriptomic and chromatin accessibility data offer a comprehensive and reliable catalogue of cell types, providing valuable insight into understand the pathogenesis of NEC across multiple molecular dimensions.

### Experimental NEC impairs proliferation and stemness in heterogenous epithelial cell populations along the villus axis

Among the altered cell types, epithelial cells displayed the most pronounced changes in chromatin accessibility and constituted the largest cell compartment in the ileum. We identified a total of 11,266 nuclei from epithelial cells in our snRNA-seq data. Initial unsupervised clustering analysis of epithelial cells identified 22 clusters. Due to the relatively small number of Paneth cells, they did not form an independent cluster. We observed that cluster 9 contained both Paneth cells (*Lyz1*^hi^) and goblet cells. Sub-clustering of cluster 9 was further performed to refine the annotation of Paneth cells (*Lyz1*^hi^ and *Defa17*^hi^). Finally, unsupervised sub-clustering of the epithelial cluster identified 7 major cell types including enterocytes (*Abcc2*, *Apob*), goblet cells (*Agr2*, *Fcgbp*, *Muc2*), Paneth cells (*Defa17*, *Lyz1*, *Spink4*), intestine stem cells (*Lgr5*), transient amplifying (TA) cells (*Hmga2*, *Top2a*, *Pcsk5*), L cells (*Pyy*, *Neurod1*), N cells (*Nts*), and enterochromaffin (EC) cells (*Chgb*, *Glis3*) (Fig. 2A-B).

**Figure 2.**
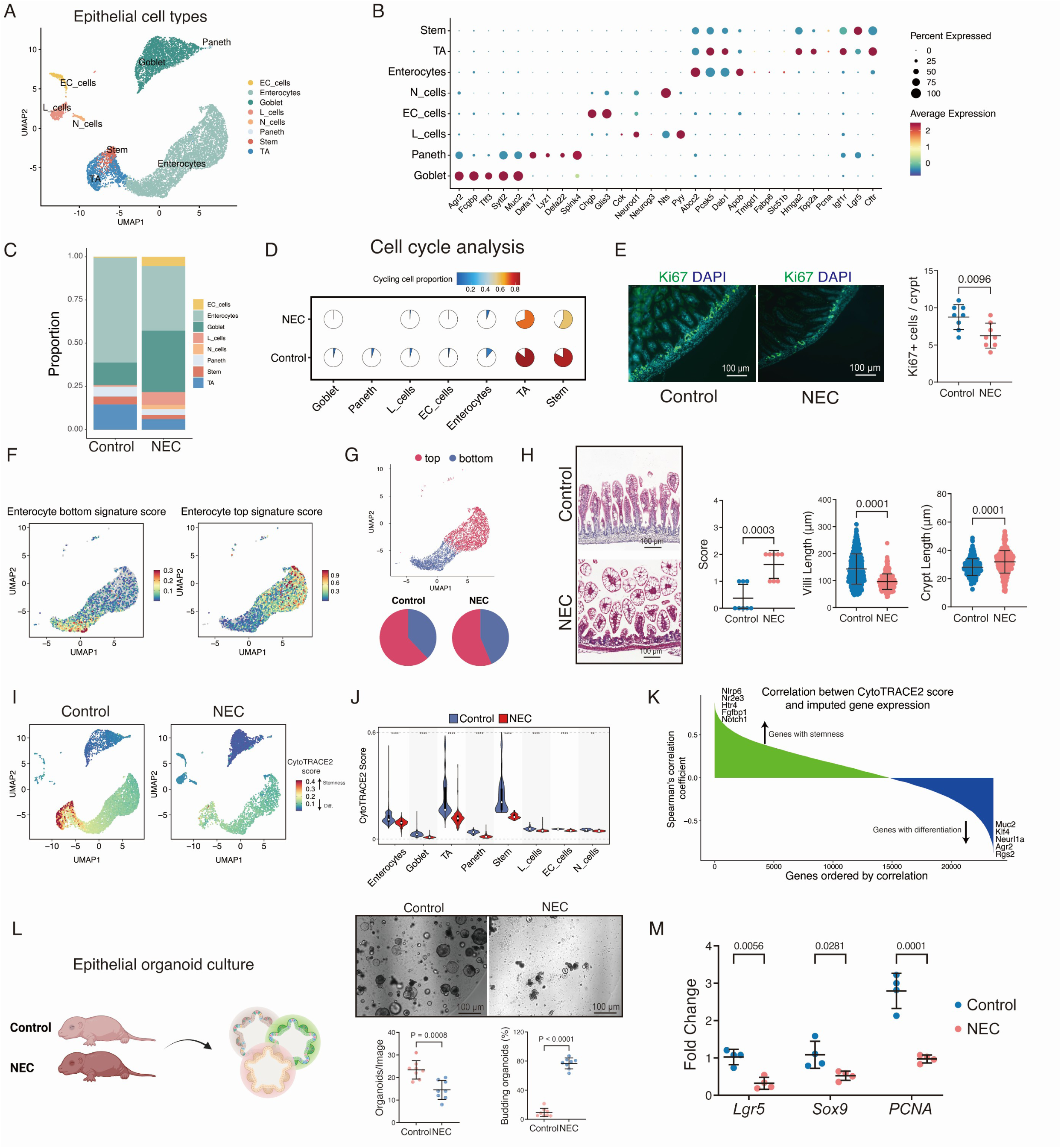
Impaired proliferation and stemness in the epithelium during experimental NEC. (A) UMAP view of epithelial cell types in snRNA-seq data. (B) Expression levels and frequencies of selected markers across epithelial cell types. (C) Barplot depicting the proportion of epithelial cell types in control and NEC group. (D) Pie chart visualizing the proportion of cycling cells of different epithelial cell subtypes from different groups. (E) Representative immunofluorescence micrographs stained for proliferation marker, Ki67, with DAPI nuclei counterstain in the ileum of control and NEC mice, and its quantification. (F) UMAP view of enterocyte bottom and top signature scores. (G) UMAP view of enterocyte category and pieplot showing proportion of top and bottom category. (H) Representative H&E-stained histomicrographs in the ileum of control and NEC mice, and the quantitative account of intestinal injury, crypt length, villi length and crypts density (10 crypts length). (I) UMAP view of CytoTRACE2 scores. (J) Violin plot showing the CytoTRACE2 scores across epithelial cell types. Significance was determined by Wilcoxon tests. (K) Spearman’s correlation between CytoTRACE2 score and imputed gene expression from MAGIC. Genes with stemness or differentiation were highlighted. (L) Representative organoid micrographs from small intestine of control and NEC mice, and quantification of percentage of organoids budded in each region of interest (ROI) and number of organoids in in each region of interest (ROI) after 5 days in cultures. (M) Relative gene expression in the control and experimental NEC mice measured by qPCR. Scale bars are shown in all the images. Experiments were repeated independently 3 times, with similar results. Each dots represented average value of each individual. Data are presented as mean ±SD and compared using student T test or one-way ANOVA with post-hoc tests as appropriate. The primary p-value is indicated directly on the graph.

Our snRNA-seq data confirmed a decrease in enterocytes, intestinal stem cells (ISCs), and transit-amplifying (TA) cells in NEC samples (Fig. 2C and Fig. S1D), with statistical significance (P < 0.05, Monte Carlo/permutation test). This aligns with the known characteristics of NEC, which include dysfunctional enterocytes and a reduced pool of intestinal progenitor cells [26]. Additionally, we observed a decreased proportion of Paneth cells in NEC compared to the control, consistent with previous studies [27, 28] (Fig. 2C and Fig. S1D). We also noted an increase in enteroendocrine cells in NEC, similar to findings in inflammatory bowel diseases (IBDs) [29] (Fig. 2C and Fig. S1D). Next, we compared the characterized proliferation status of the epithelial cell types. We observed a reduced number of cycling cells across various epithelial cell types, especially in stem cells and TA cells, during NEC (Fig. 2D). The result was validated by KI67 staining and quantification, which suggests decreased proliferation during NEC (Fig. 2E).

To characterize enterocyte differentiation programs along the villus axis, we used a gene signature [30] to cluster enterocytes into two categories: enterocyte progenitors (bottom villus-like enterocyte) and fully differentiated enterocytes (top villus-like enterocyte) (Fig. 2F-G). During NEC, we observed a decrease in top-like enterocytes, a phenomenon we refer to as “villous blunting”, confirmed this by pathological staining which revealed an increase in crypt length and a decrease in villus length, characteristic of higher NEC severity scores (Fig. 2H).

To describe the altered differentiation programs across the villus axis, we performed CytoTRACE2 [31] analysis to infer the differentiation status of all epithelial clusters. Stem cells and TA cells ranked the highest in CytoTRACE2 scores, indicating their prominent roles in maintaining stem cell characteristics and progenitor roles (Fig. 2I). We observed that overall CytoTRACE2 scores decreased across all epithelial cell types in NEC, suggesting impaired stemness in NEC epithelial cells (Fig. 2J). Furthermore, to identify genes highly correlated with cell potency and stemness, we performed MAGIC analysis and calculated the correlation between CytoTRACE2 scores and imputed gene expression from MAGIC (Fig. 2J). We identified a set of genes positively correlated with CytoTRACE2 scores, such as *Htr4*, *Nlrp6*, *Fgfbp1*, *Notch1*, and *Nr2e3* (Fig. 2K). These genes are known to play roles in pathways supporting stem cell maintenance, proliferation, and differentiation potential [32–34]. We also identified genes involved in epithelial differentiation, such as *Muc2*, *Klf4*, *Neurl1a*, *Agr2*, and *Rgs2* (Fig. 2K). To validate these changes in stemness programs, we employed an intestinal epithelial organoid model ideal for studying stem cell activity in NEC [35]. We found that intestinal epithelial organoids derived from NEC mice demonstrated a reduction in number and size, as well as reduced budding ability, compared to organoids derived from the control group (Fig. 2L). This indicates an impairment of ISC activity in NEC, validated by decreased gene expression of *Lgr5* and proliferation markers *Sox9* and *Pcna* in organoids derived from NEC (Fig. 2M).

To characterize the epigenetic changes in epithelial cell types, and their association with stemness and differentiation, we obtained a total of 11,229 nuclei from pooled epithelial cells in our snATAC-seq data, with 5,672 from the control group and 5,557 from the NEC group (Fig. S3A-B). MACS2 analysis identified 162,264 peaks, of which 34,665 were recognized as DARs. Enterocytes, goblet cells, and TA cells had the highest number of DARs. Approximately 12.3% (0.123 ±0.05) of DARs were closely associated with DEGs in their corresponding cell types. The majority of DARs were located in the intronic and distal regions of the genome, and their distribution was relatively conserved across cell types (Fig. S3C-D). Additionally, transcription factor footprinting analysis revealed transcription factors (TFs) enriched in different epithelial cell types, such as BACH1/MAFK in enterocytes, HOXA11 in TA cells, ARID5a in goblet cells, SOX6 in stem cells, PTF1A in L cells, TCF21 in Paneth cells, and ARID3b in EC cells (Fig. S3E).

Overall, these findings highlight that NEC involves cell-type-specific epigenetic reprogramming, impacting stem cell function and the differentiation capacity of epithelial cells.

### Spatial alterations in intestinal epithelial cells are revealed by MERFISH

To analyze transcriptomic changes at sub-cellular resolution across different cell types and spatial regions of the intestine [36], we applied MERFISH imaging to interrogate the spatial organization of intestinal cells identified by snRNA-seq in control and NEC conditions (Fig. 3A). A total of 495 probes (Table S2) were designed for capturing major signaling pathways or cell type genes (see *Methods*). Since NEC affects the epithelial remodeling in the intestine, we first manually annotated the anatomical regions of the intestine by assigning cells into the villus and crypt regions. Differential gene expression analysis comparing NEC villus-crypts to their respective controls revealed upregulated *Muc2*, *Lrp6*, *Myc*, *Fzd6*, and *Angptl4*, and decreased expression of *Mki67*, *Mcm5*, and *Hmgb2* in NEC in the crypt region. *Lrp5*, *Fzd5*, *Ccnd2*, and *Lrp6* genes were increased, while *Alpi* was decreased in NEC in the villus region (Fig. 3B, S4A-B). Furthermore, MERFISH revealed spatially altered cell type proportions in NEC, such as decreased enterocytes and increased SMCs in the crypt region (Fig. 3C). GO term enrichment analysis suggested spatially differentially activated pathways (Fig. 3D). For example, WNT signaling pathways, neurogenesis, and the regulation of endothelial cell proliferation were compromised in NEC in the crypt region, while pathways such as epithelial cell apoptotic process and fibroblast proliferation were activated. In the villus region, pathways related to macrophage differentiation and muscle adaptation were activated in NEC, while WNT signaling, neurogenesis, amino acid metabolic process, fluid shear stress, and thyroid hormone generation pathways were downregulated in NEC. These findings reveal that NEC induces profound spatial reorganization and functional changes in the intestine, impacting key pathways essential for epithelial integrity and homeostasis.

**Figure 3:**
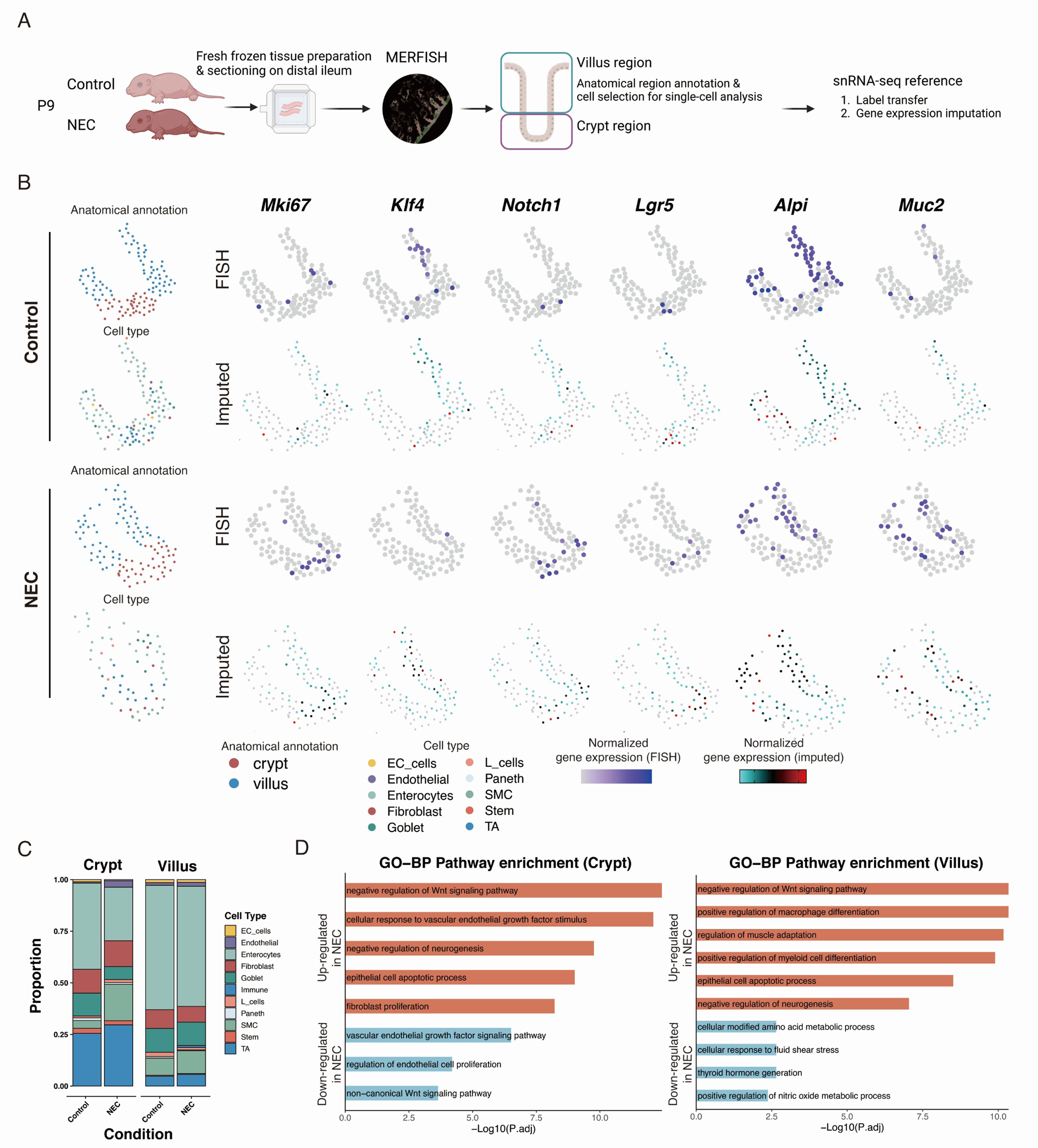
MERFISH spatial maps illustrate the alteration of intestinal epithelial cells. (A) Schematic plot of MERFISH data processing. (B) Typical regions of anatomical annotation and predicted cell types from snRNA-seq. Gene expression was shown with marker genes measured by MERFISH and imputed by the iSpatial algorithm. (C) Barplot showing the cell proportions in the crypt and villus regions in MERFISH datasets. (D) GO enrichment analysis of genes upregulated and downregulated in NEC in the crypt and villus regions.

### Integrated omics analysis revealed epigenetic regulation of epithelial developmental trajectories in the neonatal intestine during NEC

To simultaneously investigate transcriptional and epigenetic changes in NEC, we integrated snATAC-seq and snRNA-seq data using CCA to co-embed the two modalities. This approach first identifies “anchor” cells representing shared biological states in a common lower-dimensional space, then finds representative cells from one modality in the other modality. Next, we applied the scOptmatch [37] algorithm to find the optimal ATAC-RNA cell pairs, identifying 11,229 ATAC-RNA cell pairs in our dataset. This computational pairing method allows us to investigate chromatin profiling as well as gene expression per cell type (Fig. 4A). Consistent with previous studies [37], we found that chromatin accessibility at distal peaks is highly cell type-specific, even more so than gene expression, whereas promoter accessibility is relatively invariant across cell types and experimental conditions (Fig. S4C).

**Figure 4.**
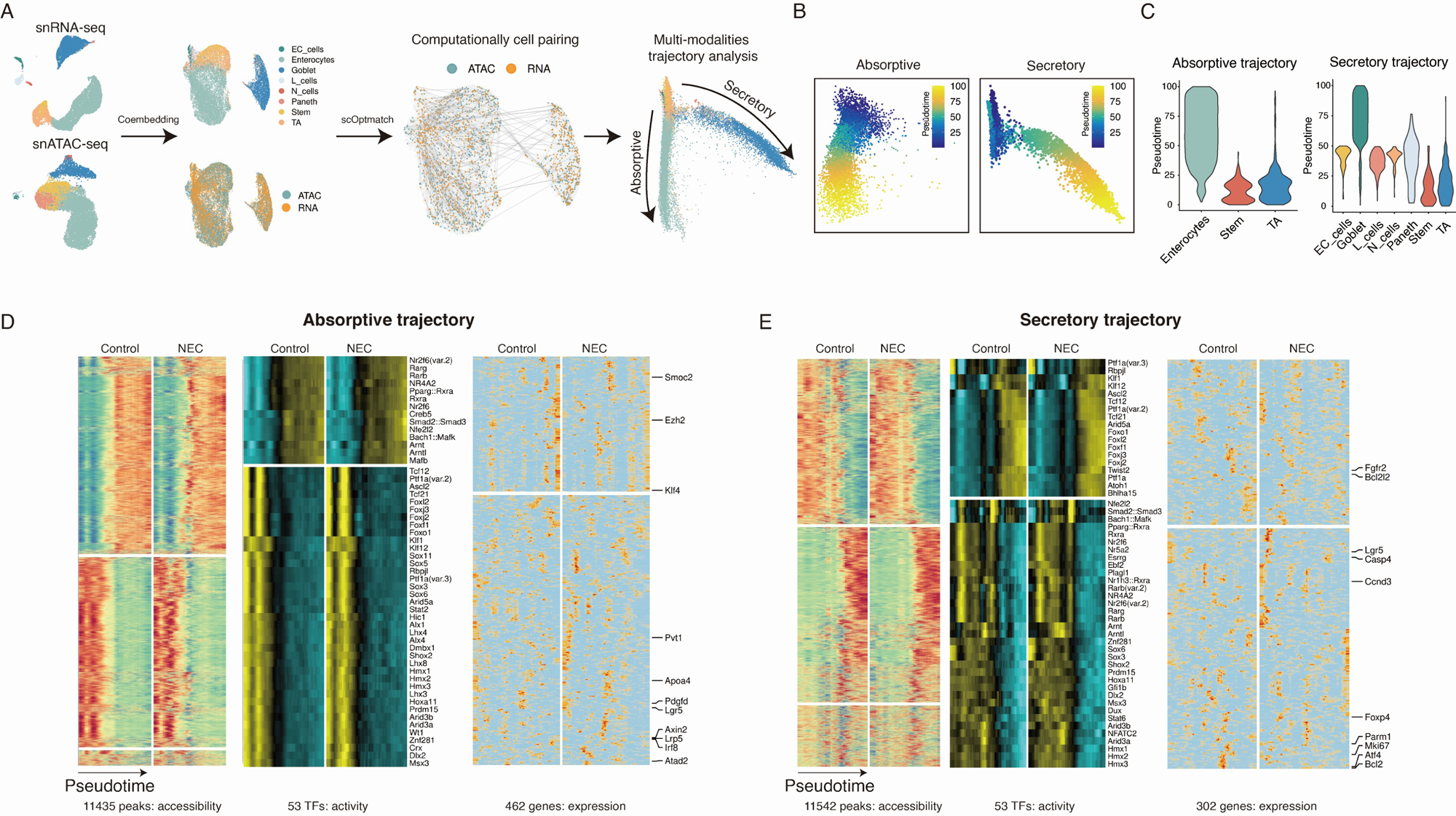
Multi-modalities trajectories analysis in epithelial cells. (A) Schematic diagram depicting integrating snRNA-seq and snATAC-seq data for multi-modalities trajectory analysis. (B) Diffusion map of the absorptive trajectory and the secretory trajectory. Cells were colored by trajectory pseudotime. (C) Violin plot showing the pseudotime in each cell type in the absorptive trajectory and the secretory trajectory. (D) Variable peaks, TFs and genes identified along the absorptive differentiation trajectory. (E) Variable peaks, TFs and genes identified along the secretory differentiation trajectory.

We created a diffusion map to explore specific developmental trajectories, identifying two primary branches rooted in stem cells and transit-amplifying (TA) cells: an absorptive branch (leading to enterocytes) and a secretory branch (leading to goblet cells, enteroendocrine cells, and Paneth cells) (Fig. 4B). The differentiation trajectory was further analyzed using pseudotime, inferred from CytoTRACE2 scores, to reveal how cells progress along each branch (Fig. 4C). This approach allowed us to map out regulatory and gene expression changes that characterize these paths.

In the absorptive branch, we identified 11,435 variable peaks representing cis-regulatory elements, capturing the regulatory dynamics that drive differentiation. Early in this trajectory, accessibility of binding motifs of TFs such as *SOX3*, *SOX6*, and *ASCL2* were found to be highly expressed, indicating their roles as master regulators that help maintain progenitor status before differentiation. This suggests that changes in these TFs in NEC could affect the availability of progenitor cells, potentially compromising the intestinal regenerative capacity. Through trajectory pseudotime analysis using tradeSeq [38], an algorithm to highlight genes with different expression patterns across control and NEC. We identified key genes involved in WNT signaling, such as *Axin2* and *Lrp5*; canonical stem cell markers, such as *Lgr5*, *Smoc2*, and *Atad2* which are crucial for maintaining stemness and initiating differentiation [32]. Additionally, the expression of *Ezh2*, a subunit of the polycomb repressive complex 2 (PRC2), suggests that epigenetic regulation through PRC2 might play a role in controlling differentiation timing, while enterocyte differentiation markers *Klf4* and *Apoa4* mark the completion of the absorptive pathway (Fig. 4D). These findings indicate that dysregulation of these pathways in NEC could alter the balance between stem cell maintenance and differentiation, contributing to the disease pathology by reducing the absorptive function of the gut epithelium.

A similar analysis of the secretory branch revealed distinct changes in cis-regulatory elements, TFs, and gene expression patterns. For instance, early enrichment of ESRRG suggests it may act as a regulator that could be negatively modulated by PRC2, possibly affecting secretory cell lineage commitment [39]. TFs such as *Atoh1*, *Foxj2*, and *Foxj3* peaked later in pseudotime, underscoring their potential roles as key regulators of goblet cell differentiation. The differential expression of genes involved in proliferation, such as *Mki67* and *Ccnd3,* and apoptosis, such as *Bcl2*, *Casp4*, and *Bcl2l2*) between control and NEC conditions (Fig. 4E) indicates that NEC disrupts the normal balance of cell survival and death along these trajectories, potentially leading to impaired epithelial renewal and barrier function.

Taking together, these findings reveal the NEC impacts on differentiation programs in the neonatal intestine, disrupting the balance of critical cell types and contributing to disease pathogenesis through impaired epithelial renewal and functionality.

### Module and differential gene analysis identify distinct pathogenic processes in NEC

We then sought to identify signaling pathways and biological processes that are changed in the identified epithelial cell types. We first performed HotSpot analysis, which clusters gene expression profiles into modules. HotSpot recognizes genes with high local autocorrelation and is robust to batch effects [40, 41]. This analysis identified 14 modules in all epithelial cells. Module analysis revealed differentially enriched genes linked to distinct functions across different cell types (Fig. 5A-B). Enteroendocrine cells showed enrichment in functions related to peptide transport, synapse transport, and axon guidance, reflecting their role in sensing ingested nutrients and regulating behavior response to feeding (Modules 1, 2, and 11). Protein glycosylation-related biological activity was enriched in goblet cells (Module 3). Paneth cells scored highest in Module 8, which overrepresented genes related to the defense response to bacteria. Modules 6 and 12 were enriched in enterocytes, aligning with their roles in lipid metabolism and sodium ion transport. Moreover, we found that the scores of Modules 5 and 6 were decreased in NEC, suggesting impaired activity in cell cycling and lipid metabolism, respectively. Conversely, several module scores were elevated in NEC, indicating pathological alterations such as glycosylation and muscle migration (Fig. 5C-D). In addition, we evaluated gene signature scores from the HARMARK gene set in our snRNA-seq data using the UCell package. We observed a reduction in E2F gene signature scores in epithelial cell types associated with NEC, particularly in stem cells and TA cells (Fig. 5E). Furthermore, DNA repair gene signature scores were also diminished in these cell types in NEC. The WNT beta-catenin pathway scores were found to be decreased in stem cells but increased in goblet cells and enterocytes, suggesting altered signaling related to the pathogenesis of NEC (Fig. 5E). Hypoxia and inflammatory response pathway scores were increased in stem cells, TA cells, goblet cells, and enterocytes highlighting the crucial role of hypoxia and inflammation in NEC pathogenesis (Fig. 5E).

**Figure 5.**
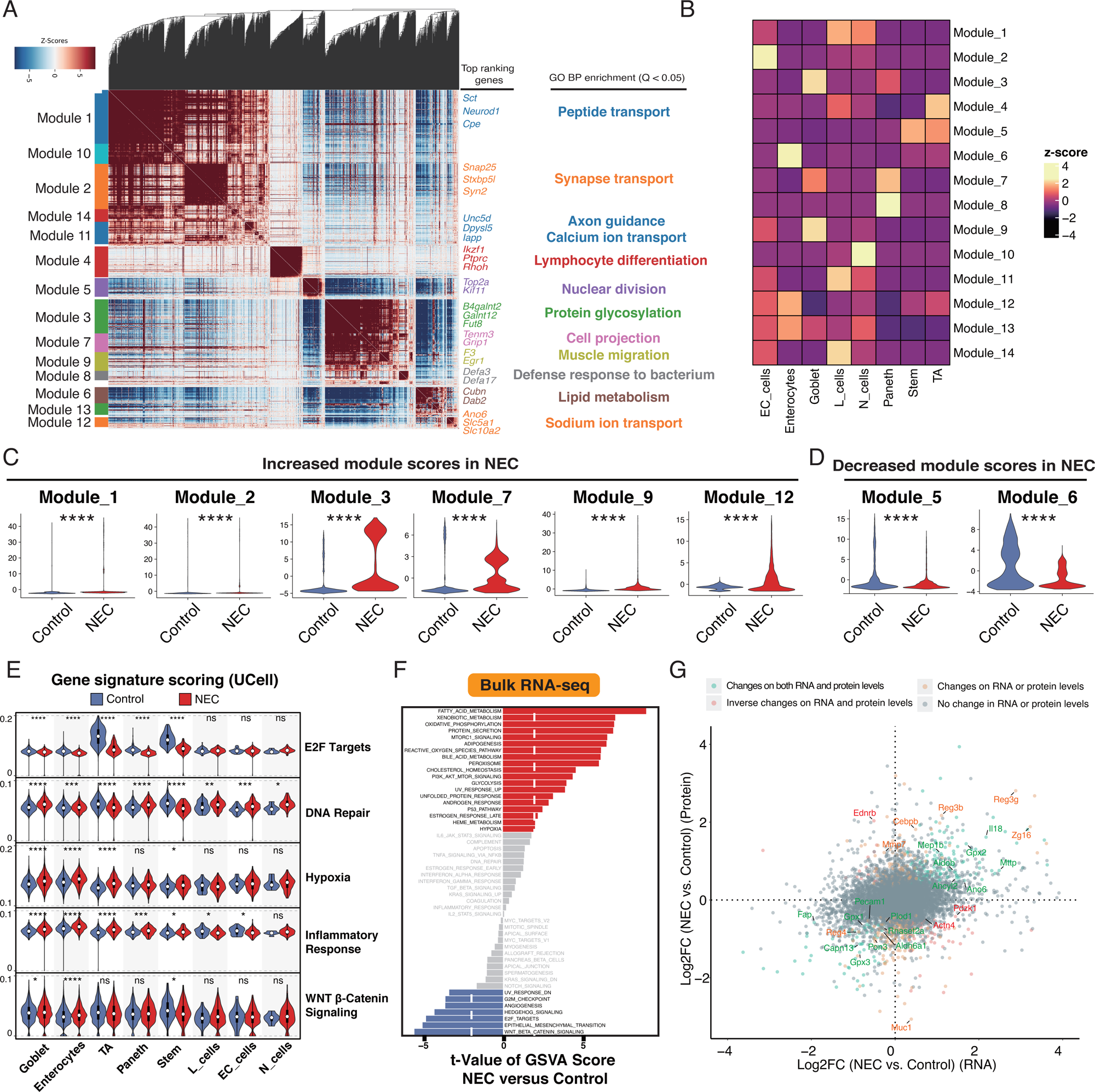
Module analysis and pathway analysis in epithelial cells. (A) The scaled gene expression level (z-score) across co-expression modules. Selected enriched genes and their associated pathways (coloured according to module) are highlighted on the right. GO, Gene Ontology. (B) The scaled aggregated expression levels (z-score) in each epithelial cell population for the co-expressing genes in each module. (C) Violin plot showing increased module scores in NEC. (D) Violin plot showing decreased module scores in NEC. (E) Violin plot showing selected gene signatures from HALLMARK gene sets enrichment scores in each epithelial cell population across conditions. Significance was determined by Wilcoxon tests. (F) Differences in pathway activity were analyzed using GSVA and t values were shown from a linear model (n = 3). (G) Scatter plot of the differentially expressed genes in the mRNA level (x-axis) and the protein level (y-axis).

To validate the completeness of our snRNA-seq data and capture alterations in NEC in both mRNA and protein content for the whole intestine, we performed bulk RNA-seq sequencing (n = 3 for each group) and mass spectrometry-driven proteomics (n = 4 for each group) for P9 mice from the NEC and control groups. From the bulk RNA-seq data, we observed an overall increase in the activity of fatty acid metabolism and hypoxia pathways, along with a decrease in the activity of E2F targets and WNT beta-catenin pathways in NEC (Fig. 5F). These findings further corroborate our snRNA-seq data. Differential gene and protein expression analysis identified 6262 differentially expressed genes and 602 proteins (See method) (Fig 5G). We observed genes such as *Il18*, *Mep1b*, *Aldob*, *Gpx2*, *Ahoyl2*, *Ano6*, *Fap*, *Pecam1*, *Capn13*, and *Aldh6a1* as having concordant changes at both RNA and protein levels. Genes such as *Ednrb*, *Pdzk1*, and *Actn4* exhibited inverse changes between RNA and protein levels. Other genes such as *Mmp7*, *Cebpb*, *Reg3b*, *Reg3g*, *Muc1*, and *Zg16* showed differences in either RNA or protein levels between NEC and control groups. These results indicate that different genes exhibit changes at both the mRNA and protein levels. The discordant changes between mRNA and protein levels suggest complex post-transcriptional regulation during the pathogenesis of NEC.

These results identify distinct functional disruptions and regulatory complexities in epithelial cell types during NEC, providing insights into key pathogenic processes and their underlying molecular mechanisms.

### Master Regulators and Progenitor Cell Dynamics in NEC-Associated Epithelial Dysfunction

To identify master regulator (MR) proteins, which determine cell fate and differentiation, we used the VIPER method [42] to infer the protein activity of transcriptional and signaling proteins from snRNA-seq data (See method). We identified 689 and 921 regulon activities in control and NEC conditions respectively. UMAP analysis revealed sharper segregation of epithelial cell populations and different segregation statuses of cell types between control and NEC (Fig. 6A). For instance, compared to UMAP reduction from snRNA-seq gene expression, Paneth cells formed a separate cluster in the VIPER analysis. Stem and TA cells co-segregated into one cluster, indicating their progenitor role. However, in NEC, stem and TA cells co-segregated with enterocytes into one larger cluster, suggesting the similarity of MR activity in these cell types. In addition, few Paneth cells were detected in NEC, clustered with goblet cells indicating shared functions of these populations. We identified highly expressed MRs in all cell types per condition (Fig. 6B). MRs such as EED and SUZ12 were upregulated in stem cells and TA cells; both are core components of PRC2, highlighting their critical role in maintaining stemness and regulating differentiation. Different highly expressed MRs were identified in NEC, with only a few overlapping across conditions, indicating distinct regulatory mechanisms in epithelial cell populations (Fig. 6C).

**Figure 6.**
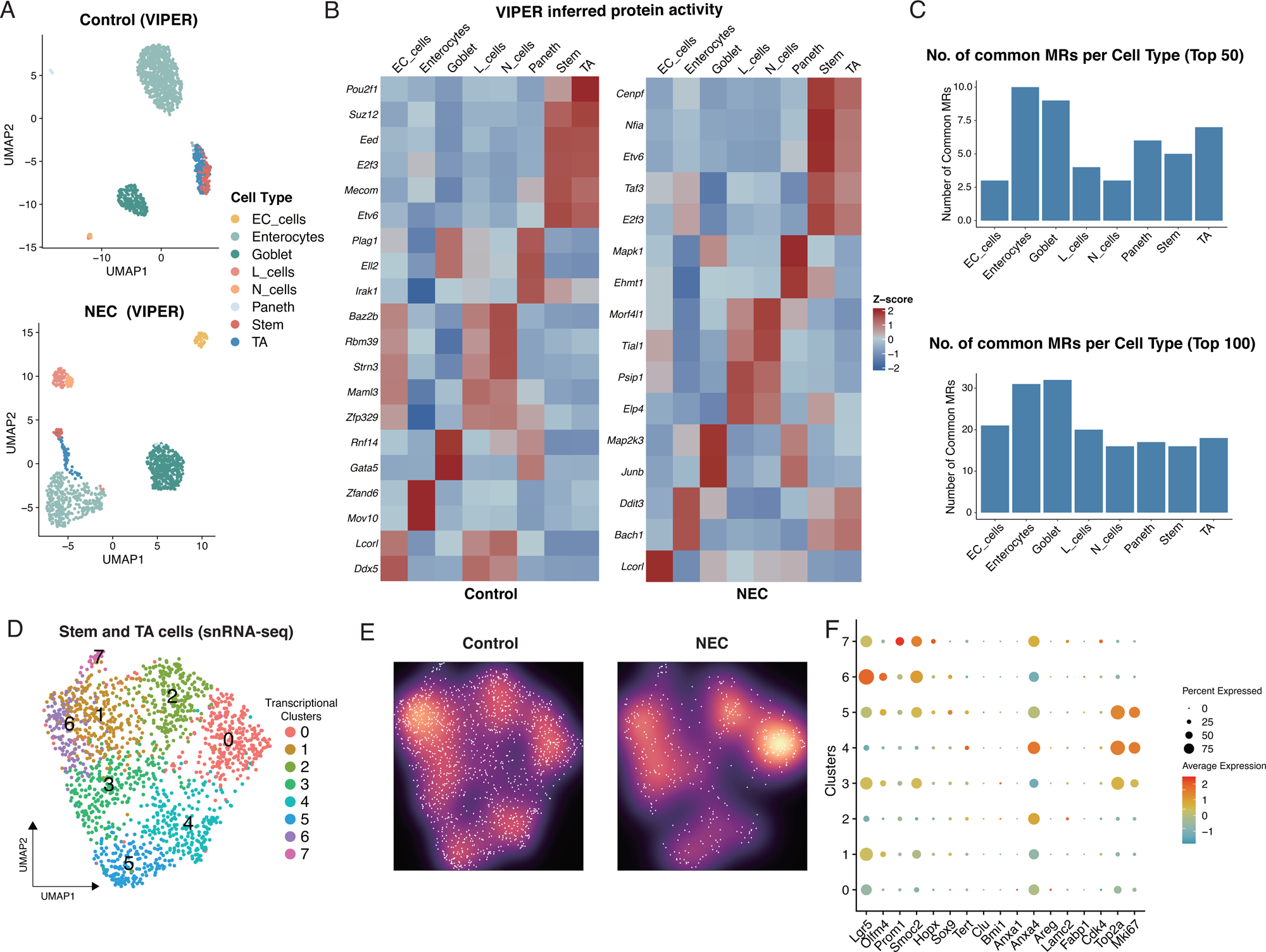
VIPER analysis revealed master regulators in epithelial cells. (A) UMAP projection of inferred protein activity from VIPER analysis in the control and NEC groups. (B) Expression level of inferred protein activity from VIPER analysis in each epithelial cell population in the control and NEC groups. (C) Barplot showing the number of common master regulators (MRs) in top 50 (top) or top 100 (bottom) MRs in each cell type between control and NEC group. (D) UMAP view of clusters in intestinal stem cells and TA cells. (E) Galaxy plots depicting cell density in UMAP for cells from the control and NEC groups. Cooler colors indicate low density, and warmer colors indicate high density. (F) Expression levels and frequencies of selected markers across clusters.

Building on these findings, we explored how shifts in MR activity influence the composition and transcriptional landscape of progenitor cell populations, with a particular focus on stem and TA cells, which are essential for maintaining epithelial homeostasis and regeneration. Through snRNA-seq data analysis, we identified 1,662 stem and TA cells, which were further categorized into seven distinct transcriptional clusters, underscoring their heterogeneity (Fig. 6D). The distribution of these clusters varied between NEC and control, indicating differences in the progenitor population (Fig. 6E). Genes related to stemness were differentially enriched in clusters. *Lgr5* was highly expressed in clusters C1, C3, and C6, while *Anxa4* was highly expressed in C5 (Fig 6F). *Top2a* and *Mki67* were preferentially expressed in clusters C3, C4, and C5. However, stemness markers such as *Tert*, *Clu*, and *Bmi1* were rarely expressed in NEC and controls, suggesting that they may not be critical stemness regulators in the neonatal intestine (Fig. 6F).

Overall, these findings reveal significant disruptions in master regulator activity and progenitor cell composition in NEC, underscoring the complexity of epithelial dysfunction and highlighting potential targets for therapeutic intervention.

### Reduction of EZH2 expression in the epithelial progenitor cells leads to the dysregulation of ISCs and severe intestinal injury in experimental NEC

In light of our previous study demonstrating the dysregulated Lgr5 ISC expression in experimental NEC [12], we next aimed to investigate how alterations in the PRC2-EZH2 epigenetic pathway regulate ISCs and contribute to NEC pathogenesis. NEC mice exhibited decreased *Lgr5* gene expression during NEC progression (Fig. 7A), with *Ezh2* showing a similar expression pattern (Fig. 7B). Concurrently, low expressions of Ezh2 and H3K27Me3 were observed in the crypt base, where Lgr5+ ISCs reside (Fig. 7C).

**Figure 7.**
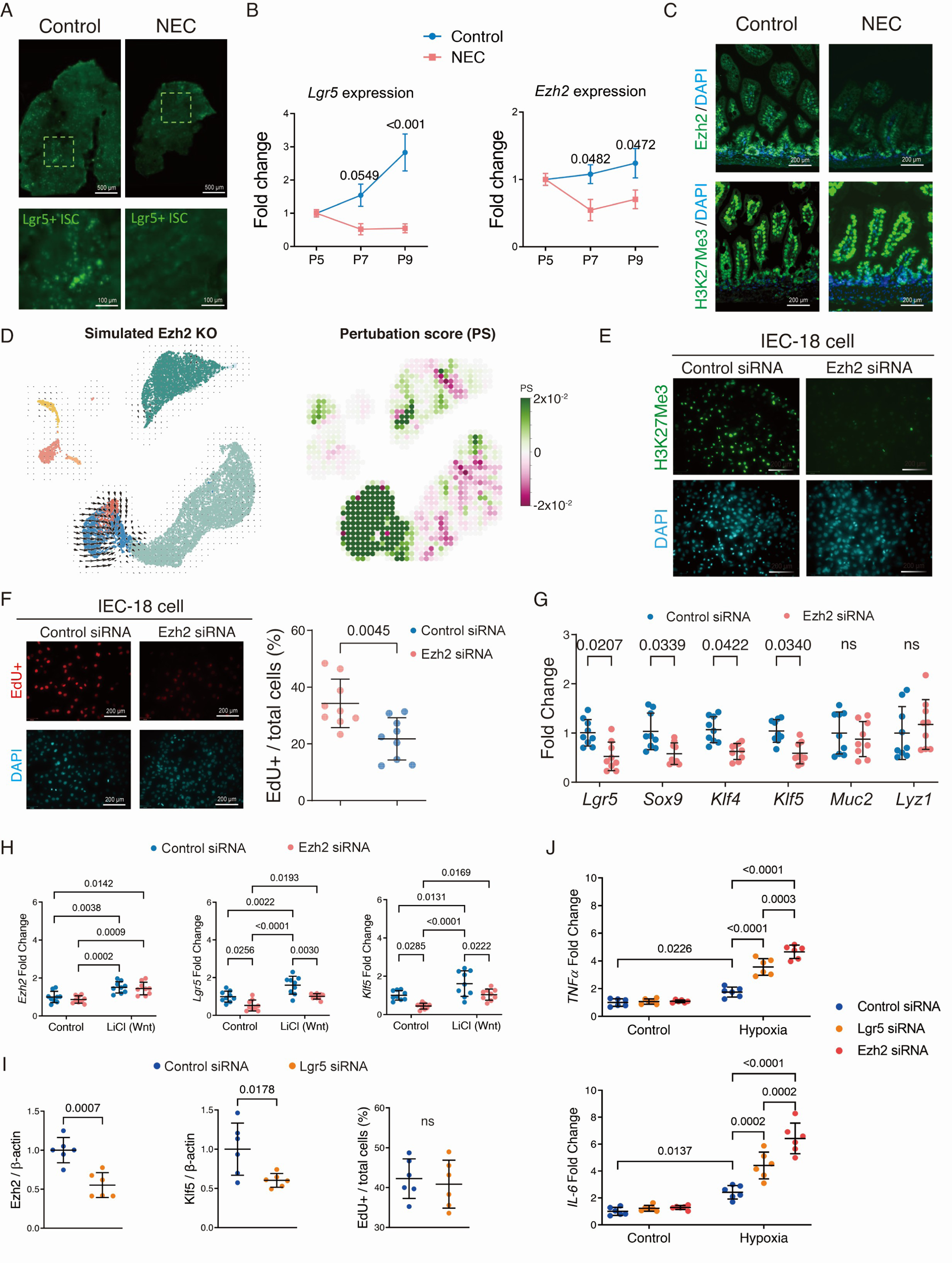
Ezh2 is directly involved in maintaining Lgr5+ ISCs, which may influence intestinal proliferative and inflammatory responses *in vitro*. (A) Representative fluorescence micrographs of the intestine, obtained from control and NEC Lgr5+ GFP mice. Green dots indicate the presence of LGR5+ ISCs. (B) Relative expression of *Lgr5* and *Ezh2* between postnatal days 5 to 9 in mice induced with experimental NEC compared to breastfed controls. (C) Representative immunofluorescence micrographs stained for Ezh2 and H3K27Me3 in mice with experimental NEC, with DAPI nuclei counterstaining. (D) CellOracle algorithm to perform an *in silico* knockdown of *Ezh2* in epithelial cells. (E) Representative immunofluorescence micrographs of H3K27me3 in the Ezh2 siRNA-treated IEC-18 cells, with DAPI nuclei counterstaining. (F) Representative immunofluorescence micrographs and quantification of EdU+ proliferating intestinal epithelial cells in *Ezh2* siRNA-treated IEC-18 cells, with DAPI nuclei counterstaining. (G) Relative gene expression in IEC-18 cells treated with Ezh2 siRNA compared to the control. (H) Relative gene expression in IEC-18 cells treated with Ezh2 siRNA in the presence of the Wnt pathway activator, Lithium Chloride (LiCl), compared to the control. (I) Quantification of EZH2 and KLF5 protein expression and quantification of EdU+ proliferating cells in the *Lgr5* siRNA-treated IEC-18 group. (J) Relative expression of TNFα and IL-6 inflammatory cytokines in IEC-18 cells administered with either *Ezh2* or *Lgr5* siRNA in response to inflammation induced by hypoxia. Scale bars are shown in all the images. Experiments were repeated independently 3 times, yielding similar results. Each dot represents the average value for each individual. Data are presented as mean ± SD and compared using student T-test or one-way ANOVA with post-hoc tests as appropriate. The primary p-value is indicated directly on the graph.

To study the effect of Ezh2 perturbation on NEC epithelial injury, we used the CellOracle algorithm to perform an *in-silico* knockdown-(KD) of *Ezh2* in epithelial cells. The simulated *Ezh2* KD primarily predicted an effect on ISCs and TA cells, promoting their differentiation while inhibiting the maturation of enterocytes. This could explain the decreased stemness and enterocyte maturation observed in NEC (Fig. 7D). To further validate these findings, we then silenced *Ezh2* using siRNA in IEC-18 cells, which resulted in reduced EZH2 protein levels without altering mRNA levels (Fig. S4D and F). *Ezh2* KD led to a reduction of H3K27Me3 levels (Fig. 7E) and decreased proliferation of EdU-stained IEC-18 cells (Fig. 7F). Stemness and differentiation genes, including *Lgr5*, *Sox9*, *Klf4*, and *Klf5*, were downregulated after *Ezh2* KD, whereas secretory linage maker *Muc2* and *Lyz1* genes were unaffected. This suggests that *Ezh2* influences intestine progenitor stemness and enterocyte differentiation, while other cell types, such as goblet cells and Paneth cells, are not impacted (Fig. 7G).

Since ISC stemness is activated by the Wnt pathway, we investigated if upregulating Wnt signaling by an agonist, LiCl, influences *Ezh2* activity. We found that *Ezh2*, *Lgr5*, and *Klf5*, are upregulated upon exposure to LiCl (Figure 7H). Further, we found that the activation of *Lgr5* and *Klf5* is inhibited by *Ezh2* siRNA despite LiCl exposure, indicating that *Ezh2* acts possibly upstream of *Lgr5* within the Wnt pathway (Fig. 7H). Silencing *Lgr5* in IEC-18 cells resulted in a reduction of *Ezh2* and *Klf5* expression, suggesting a possible feedback loop from *Lgr5* to *Ezh2* expression (Fig. 7I and S4E-F). Since signaling by proinflammatory cytokines IL-6 and TNFα in the epithelium, driven by hypoxic conditions, is a hallmark of NEC pathogenesis [43], we modeled injury in IEC-18 cells *in vitro* using intermittent hypoxia to simulate these conditions. We confirmed that hypoxia induced upregulation of pro-inflammatory cytokines IL-6 and TNFα in IEC-18 cells, contributing to injury. However, silencing either *Lgr5* or *Ezh2* resulted in further increased *Il-6* and *Tnfα* mRNA expression in response to hypoxia-induced intestinal injury, with the *Ezh2* siRNA group showing the highest levels of expression (Fig. 7J). These findings suggest that *Ezh2* is directly involved in maintaining LGR5+ ISCs, which may influence intestinal proliferative and inflammatory responses *in vitro*.

To validate the *in vitro* findings, *Ezh2* conditional knockdown was induced in LGR5+ ISCs in an *in vivo* mouse model from P3 to 5 before the experimental NEC induction from P5 to 9 (Fig. 8A). At P5, *Ezh2* knockdown mice exhibited alterations in epithelial morphology and total mortality (Fig. 8B-C). A decrease in *Lgr5* expression was observed in knockdown mice compared to controls accompanied by a reduction in the number of Ki67-marked proliferating epithelial cells (Fig. 8D-F). In addition, intestinal organoids derived from these mice exhibited similar phenotypes to those derived from NEC mice, with more budding and smaller size (Fig. 8G-H).

**Figure 8.**
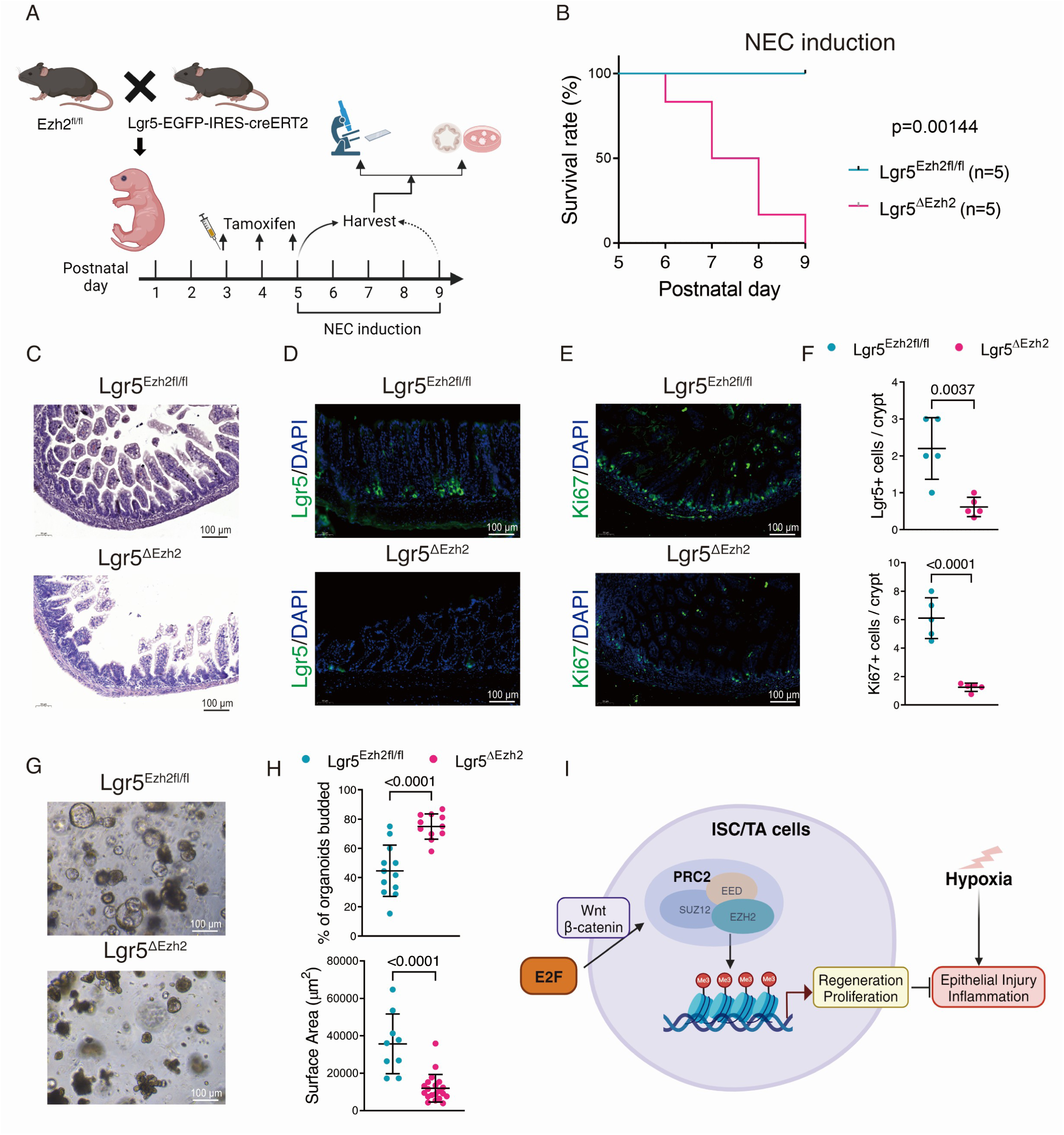
Reduction in *Ezh2* expression in the crypts leads to dysregulation of ISCs and severe intestinal injury in experimental NEC. (A) Schematic illustration of animal experimental design. (B) Survival curve of the Lgr5^Ezh2fl/fl^ and Lgr5**^Δ^**^Ezh2^ mice during NEC induction. Each group has 5 mice. (C) Representative H&E-stained histomicrographs of the ileum from Lgr5^Ezh2fl/fl^ and Lgr5**^Δ^**^Ezh2^ mice. (D) Immunofluorescence staining of LGR5+ ISCs in both Lgr5^Ezh2fl/fl^ and Lgr5**^Δ^**^Ezh2^ mice groups. (E) Immunofluorescence staining of Ki67+ cells in both Lgr5^Ezh2fl/fl^ and Lgr5**^Δ^**^Ezh2^ mice groups. (F) Quantification of LGR5+ ISCs and proliferative Ki67+ cells per crypt. (G) Organoids after 5 days in cultures for both Lgr5^Ezh2fl/fl^ and Lgr5**^Δ^**^Ezh2^ mice groups. (H) Quantification of percentage of organoids budded in each region of interest (ROI) and surface area of organoids in each region of interest (ROI) image. (I) Schematic illustration of research findings in the experimental NEC model. Experiments were repeated independently 3 times, yielding similar results. Each dot represents the average value for each individual. Data are presented as mean ±SD and compared using student T-test or one-way ANOVA with post-hoc tests as appropriate. The primary p-value is indicated directly on the graph.

These results confirm that *Ezh2* is directly involved in regulating LGR5+ ISC and maintaining intestinal homeostasis. Loss of *Ezh2* leads to a reduced regenerative response to intestinal injury and inflammation (Fig. 8I).

## Discussion

We utilized a unique mutli-omics approach combining bulk RNA-seq, snRNA-seq, snATAC-seq, and MERFISH to map and describe the development of the intestinal epithelium in the neonatal mouse intestine. Our transcriptomic and chromatin accessibility data provided a robust foundation for comprehending the pathogenesis of NEC across multiple modalities. We observed significant reductions in epithelial cell proliferation and stemness markers, particularly affecting enterocytes, ISCs, and TA cells in our experimental NEC model. These changes were accompanied by dysregulation in chromatin accessibility patterns, highlighting key regulatory elements associated with differentially expressed genes. Specifically, our results implicated Ezh2 - mediated epigenetic mechanisms in ISC maintenance and intestinal regeneration, underscoring their role in NEC pathogenesis. This multi-modal omics approach offers detailed insights into NEC pathophysiology, emphasizing epithelial cell dysfunction and regulatory network alterations that contribute to intestinal injury and impaired regeneration.

The pathophysiology of NEC is complex and not yet fully understood. It involves complex regulatory networks affecting the transcriptome across different cell types of the epithelium. Impaired reconstitution of the mucosa was observed in NEC patients, aggravating bowel necrosis [44], which may lead to mucosal atrophy and bowel shortening [45]. Clinical studies have shown that in acute NEC, there is an increase in proliferative cells in areas with both severe and moderate epithelial damage. This proliferative zone is not limited to the small intestinal crypts but extends to the villi [46]. However, this proliferative response is insufficient to rapidly reverse the severe loss of mucosal lining during NEC [47]. By integrating MERFISH data with snRNA-seq, we mapped differential gene expression across the villus and crypt regions in both control and NEC conditions. In NEC-affected tissues, the crypt region showed upregulation of genes such as *Muc2*, *Lrp6*, and *Myc*, indicative of increased mucus production and cell proliferation. However, the concurrent decrease in *Mki67* and *Hmgb2* expression suggests a compromised capacity for epithelial regeneration, contributing to the impaired mucosal reconstitution observed in NEC patients. Additionally, we observed altered cell type proportions in NEC, with a notable decrease in enterocytes and an increase in smooth muscle cells within the crypts. This disrupted epithelial-mesenchymal balance likely exacerbates the pathological remodeling of the intestinal architecture, a hallmark of NEC. In the villus region, the downregulation of *Alpi* and related pathways points to compromised absorptive function in NEC-affected infants.

Our previous research demonstrated that exposure to NEC risk factors—such as hypoxia, formula feeding, and bacterial dysbiosis—leads to reduced *Lgr5* expression in the intestinal epithelium, resulting in impaired regeneration [48]. Moreover, studies have shown that LGR5+ ISCs in both NEC mice and patients exhibit reduced proliferation and increased apoptosis [12, 49]. Our sequencing analysis further revealed that NEC is characterized by impaired epithelial proliferation and reduced stemness, with distinct variations in chromatin accessibility across different cell types. Using lineage tracing techniques, we observed in mice that following NEC injury, the gut epithelium undergoes full recovery, driven by LGR5+ ISCs. However, the source of cells that replace LGR5+ ISCs lost during active NEC injury, as well as their contribution to subsequent epithelial regeneration, remain largely unexplored. To investigate the loss of LGR5+ ISCs, various studies have revealed the role of quiescent ISCs. These cells, often characterized by markers such as *Bmi1* and *Hopx*, become activated in response to injury or the loss of active stem cell populations like LGR5+ cells [50]. Lineage tracing experiments have shown that quiescent ISCs, though resistant to stress, can self-renew and give rise to all lineages of the intestinal epithelium [51, 52]. However, stemness markers such as *Hopx* and *Bmi1* were rarely expressed in our neonatal period data, suggesting that these markers may not be critical regulators of stemness in this scenario. Research has shown that Clu+ revival ISCs are resistant to intestinal injury and likely play an important role in cellular renewal following near-complete epithelial loss [53]. However, we did not observe an influx replacement from Clu+ revival ISCs, suggesting that these revival stem cells may not be the primary source of neonatal NEC.

The ISC gives rise to progenitors that subsequently differentiate into absorptive and secretory cells. Absorptive progenitors differentiate into enterocytes, which constitute the majority of the epithelium and are responsible for nutrient absorption. Secretory progenitors differentiate into goblet cells, tuft cells, enteroendocrine cells, and Paneth cells, all of which produce mucin, hormones, and antibacterial agents [54]. All differentiated cells, except Paneth cells, migrate upwards along the crypt-villus axis and eventually slough off into the lumen, in contrast, Paneth cells move downwards towards the crypt and are interspersed among crypt-based ISCs [55]. In the crypts, Paneth cells and the surrounding mesenchyme/myofibroblasts constitute a critical niche environment. They generate several secretory and/or membrane-bound ligands that tightly regulate ISC proliferation, maintenance, and differentiation [56, 57]. Mesenchymal cells play a crucial role in Paneth cell differentiation within niches during early postnatal development. Immature Paneth cells exhibit reduced Wnt secretion, contributing to instability in ISC niches and impaired regeneration in NEC [58]. In this study, Paneth cells and epithelial progenitors exhibited distinct clustering patterns when compared to controls. In NEC, although only a small number of Paneth cells were detected, they clustered with goblet cells, suggesting functional similarities between these populations. This finding aligns with the observation that Paneth cells and goblet cells share common regulatory pathways and transcription factors involved in their differentiation [59, 60]. NEC epithelial cells demonstrated unique MR profiles, highlighting distinct regulatory mechanisms in epithelial responses to disease states. Interestingly, stem and TA cells co-segregated into one cluster, indicating their progenitor roles. However, under disease conditions, stem and TA cells co-segregated with enterocytes into a larger cluster, suggesting similarities in MR activity among these cell types. This is supported by the observation that enterocyte progenitors can dedifferentiate to replace lost LGR5+ intestinal stem cells [61], suggesting that various progenitor populations can regain stemness following LGR5 reduction in NEC. Notably, spatial transcriptomics visualized alterations in cell types, with the proportion of enterocytes decreasing in the crypts but not in the villi, indicating that villus blunting affects all cell types rather than specific ones. However, remodeling occurs primarily in the crypt area. More recently, isthmus progenitor cells have been found to contribute to homeostatic cellular turnover and support regeneration following intestinal injury [42]. FGFBP1+ upper crypt cells generate progeny that propagate bi-directionally along the crypt-villus axis and serve as a source of LGR5+ cells in the crypt base [62]. These studies, together with ours, suggest an alternative model of intestinal epithelial cell organization, indicating that stemness potential is not restricted solely to the crypt regions but rather includes other drivers of intestinal regeneration.

Both the heterogeneity of epithelial progenitors and the investigation of MRs highlighted the critical role of EZH2 and its downstream H3K27Me3 targets during NEC. The reduced *Ezh2* activity was accompanied by reduced *Lgr5* expression, decreased ISC proliferation, and increased expression of inflammatory cytokines [48]. Our *in vitro* RNA interference experiments showed that EZH2 depletion decreases *Lgr5* expression, enhances inflammatory cytokine expression, and diminishes proliferation. Similarly, *in vivo* depletion of EZH2 in LGR5+ ISCs resulted in reduced LGR5+ ISCs, fewer proliferating cells, and exacerbation of the severity of the intestinal injury. While EZH2 is traditionally associated with transcriptional repression through its role in PRC2, its influence on Lgr5 expression may occur indirectly by modulating transcriptional programs critical for ISC maintenance. Further studies are required to identify the specific downstream targets of EZH2 whose dysregulation contributes to NEC pathology. Nonetheless, our findings replicate NEC-like injury characteristics and underscore the role of EZH2 depletion in exacerbating inflammation and compromising ISC function in NEC. These align with previous studies highlighting *Ezh2* and its downstream role in the Wnt pathway in ISC maintenance and intestinal epithelial regeneration [63]. In the intestine, Wnt ligands released by niche components activate ISCs and epithelial surface receptors, driving nuclear translocation of β-CATENIN and TCF4-mediated expression of Wnt target genes including *Lgr5* [64–66]. EZH2 facilitates transcriptional activation of these β-catenin target genes through PAF-mediated recruitment to the TCF4 complex, promoting intestinal homeostasis and regeneration [67]. Reduced *Ezh2* expression in NEC likely compromises β-catenin/TCF4 complex formation in ISCs, thereby suppressing *Lgr5* and other proliferation-promoting genes.

*Ezh2* is expressed globally across all cell types in the intestine [68], aberrant intestinal epithelial differentiation due to Ezh2 misregulation may disrupt this balance and subsequently drive injury. NOTCH signaling prevents ISCs from differentiating into the secretory lineage [13]. Activation of the NOTCH pathway increases the expression of Hes1 and Hes5, which inhibit the transcription of the Atoh1 factor, thereby blocking secretory lineage commitment [69–71]. A study demonstrated that inhibiting PRC2 through *Eed* knockdown led to increased *Atoh1* expression in intestinal crypt cells independently of NOTCH signaling, due to enhanced H3K27Me3 deposition at *Atoh1* promoters [13]. This suggests that the loss of PRC2 activity results in increased secretory lineage differentiation in progenitors. Interestingly, despite the expected shift towards secretory lineage differentiation, our study did not observe increased proportions of goblet cells or Paneth cells following *in vitro* Ezh2 knockdown. This discrepancy may be due to the overall knockout of Ezh2 and compensatory pathways that limit secretory cell differentiation. Indeed, Ezh2 knockdown specifically in absorptive cell lineages resulted in a marked decrease in gut barrier integrity, increased colonic permeability, and exacerbated morphological injury compared to controls [72]. In our *in vivo* experiments, immunofluorescence staining showed decreased EZH2 and H3K27Me3 in intestinal crypts housing LGR5+ ISCs, but this does not exclusively confirm effects specific to LGR5+ ISCs, as other cells are present in the crypts. Additionally, although our *Ezh2* knockdown mouse model effectively eliminates EZH2 in LGR5+ ISCs, it does not fully suppress downstream H3K27Me3 activity due to potential compensation by EZH1 and residual PRC2 components [14]. Consequently, these findings may not fully elucidate the complete downstream effects of *Ezh2*-induced transcriptional repression via histone trimethylation.

This study has certain limitations that should be acknowledged. Firstly, the findings are based on a single mouse model of NEC, which may not fully capture the heterogeneity of human NEC cases. Additionally, the sample size in each experimental group may limit the generalizability of the results, indicating that larger cohorts and validation in different models or human samples could strengthen the robustness of the conclusions. Prematurity is a pivotal factor in NEC development, contributing to critical issues such as impaired intestinal vasculature, immature immune responses, diminished regenerative capacity, underdeveloped enteric nervous system, and compromised gut motility. These factors collectively increase susceptibility to NEC, emphasizing the complex interplay between developmental immaturity and gastrointestinal vulnerabilities in its pathogenesis. Furthermore, despite utilizing advanced single-cell sequencing techniques (snRNA-seq and snATAC-seq), the study is constrained by the resolution of these methods, potentially missing rare cell populations or subtle epigenetic changes. Enhancements in sequencing depth and bioinformatics analysis are needed to improve the detection of low-abundance cell types and regulatory elements in future studies.

In summary, we observed significant reductions in epithelial cell proliferation and stemness marker expression in our experimental NEC model, accompanied by dysregulated chromatin accessibility patterns. This study highlights key regulatory elements associated with differentially expressed genes and establishes a detailed atlas that delineates molecular and cellular changes underlying NEC. These findings lay the groundwork for future more in-depth mechanistic investigations and tailored therapeutic interventions.

## Methods

### Mice

All animal experiments were approved by the Animal Care Committee at The Hospital for Sick Children (protocol no. 58119). The NEC mouse model was constructed according to methods described in previous studies [12]. Five-day-old C57BL/6 mice were separated from their mothers to prevent breastfeeding and NEC was induced by exposure to hypoxia (5% O_2_ for 10 minutes, 4 times a day), gavage feeding of formula milk (15 g Similac + 75 mL Esbilac, 50 μl/g, 4 times a day), and oral administration of lipopolysaccharide (LPS, Sigma #L2880) (4 mg/kg/day on postnatal days 6 and 8) from postnatal days 5 to 9 [12]. A group of five-day-old mouse pups that were fed by their mothers without NEC induction served as the control group. On postnatal day 9, pups from both groups were sacrificed, and their intestinal tissues were harvested for analysis. Pups were randomly assigned to each of the experimental groups to eliminate potential within-litter effects.

*Lgr5-EGFP-IRES-creERT2* mice were obtained from Jackson Laboratory (Sacramento, CA), enabling the visualization of LGR5+ ISC expression by genetically fusing the green fluorescence protein (GFP) with LGR5+ ISC [73]. Ezh2^fl/fl^ mice were obtained from P.D.O laboratory, Toronto, and established as previously reported [74, 75]. Ezh2^fl/fl^ mice were cross-bred with *Lgr5-EGFP-IRES-creERT2* mice to generate *Lgr5-EGFP; Ezh2^-/-^*mice. Cre recombinase was activated by the administration of 1 mg tamoxifen (Invitrogen) via gavage or intraperitoneal injection for 2 or 3 days. Tamoxifen was dissolved in corn-oil (Sigma) to create a stock concentration of 10 mg/ml.

### Histological staining and assessment

Mice ileal tissues embedded in paraffin were cross-sectioned (5µm) and stained with hematoxylin and eosin. Histological sections were assessed by three blinded investigators following an established histopathological scoring system [76]: grade 0 = no damage; grade 1 = epithelial cell lifting or separation; grade 2 = sloughing of epithelial cells to the mid-crypt level; grade 3 = necrosis of the entire crypt; grade 4 = transmural necrosis., as grade 0 = no damage; grade 1 = epithelial cell lifting or separation; grade 2 = sloughing of epithelial cells to the mid-crypts level; grade 3 = necrosis of the entire crypts; grade 4 = transmural necrosis.

### Immunostaining

Sections of ileal tissue were incubated with primary antibodies diluted 1 in 500 (Table S1) overnight at 4 °C. For immunofluorescence staining, sections were then incubated with secondary antibodies diluted 1 in 1000 (Table S1) and DAPI for visualization of cell nuclei (Vector Laboratories, Burlington, ON) at room temperature for 1 hour. Slides were imaged using a Nikon TE-2000 digital microscope equipped with a Hamamatsu C4742-80-12AG camera. Three blinded investigators counted the number of positively labeled cells from at least ten crypts of the intestine and five images from each subject.

### Gene quantification

RNA was isolated from ileal tissue using TRIzol (Invitrogen). Total RNA (1 µg) was reverse transcribed using qScript cDNA SuperMix (Quanta Biosciences, Gaithersburg). SYBR green-based quantitative polymerase chain reaction (qPCR) was performed using a CFX384 C1000 Thermal Cycler (Bio-Rad) and Advanced qPCR MasterMix (Wisent, Quebec) according to the manufacturer’s protocol, and primers listed in Table S1. Data were analyzed using CFX Manager 3.1 (Bio-Rad). Results are based on three independent experiments performed in triplicate. Expression levels were calculated by the ΔΔCt method and normalized to the reference housekeeping gene *glyceraldehyde 3-phosphate dehydrogenase (Gapdh)* [12].

### Protein quantification

Protein expression was quantified using immunoblotting analysis as previously described [12]. The proteins were separated and transferred to a membrane, then probed with primary antibodies (1:500) overnight at 4 °C, followed by secondary antibodies (1:1000) at room temperature for 1 hour. Immuno-positive bands were detected using an ECL Plus kit (Invitrogen, Carlsbad, CA). Band intensities were determined using an Odyssey scanner (LI-COR Biosciences, Lincoln, NE). Densitometry ratios were calculated relative to the levels of the loading control.

### Intestinal organoids

Intestinal organoids were cultured according to protocols previously described [35]. Terminal ileal tissues were harvested and cut into 1-2 mm small segments. Intestinal crypts were isolated by digestion with Gentle Cell Dissociation Reagent (StemCell Technologies, Cambridge, MA) for 15 minutes and then pelleted by centrifugation. Crypts were then resuspended in Matrigel (Corning, New York) and transferred into 24-well plates. After polymerization, the mouse IntestiCult organoid growth medium (StemCell Technologies, Cambridge, MA) supplemented with penicillin-streptomycin (100 U/mL) was overlaid on the gel in each well. Organoids were maintained in a 37°C and 5% CO2 incubator with the culture medium replaced every 48 hours. Organoids were imaged daily, and their surface area was calculated using ImageJ software 1.53e.

### IEC-18 cells

The rat intestinal epithelial cell line (IEC-18, ATCC, Manassas, VA) was cultured, as previously reported [65], and maintained in DMEM medium (Gibco) supplemented with 10% FBS and 1% Pen-strep. IEC-18 cells were transfected with established Ezh2 or Lgr5 siRNA (Qiagen, Lafayette, CO) dissolved in Lipofectamine 3000 Transfection Reagent at a concentration of 10nM (Invitrogen, Carlsbad, CA). A group of cells were also administered a negative control vector at an identical concentration to the control siRNA group. 24 hours post-transfection, cells administered with Ezh2 siRNA or control cells were analyzed. Following 24h, all cell groups were also exposed to hypoxia (5% O_2_) to induce injury and inflammation [77]. Click-iT Edu kit (Invitrogen,) was used to study the cell proliferation.

### Sample preparation for snRNA-seq

The nuclei suspension was loaded onto the Chromium Next GEM Chip M (10X Genomics, Cat. No. 1000349), and single-cell gel beads were generated in the Gel Bead-in-emulsion (GEMs) (10X Genomics, Cat. No. 1000350) using the Chromium X system, following the manufacturer’s recommendations. Captured nuclei in the single-cell gel beads were lysed, and the released RNA was barcoded by reverse transcription in individual GEMs. Barcoded, full-length cDNA was synthesized, and libraries were constructed. All libraries were sequenced in paired-end mode (PE150) for 150 bp reads on a NovaSeq 6000 platform (Illumina).

### Sample preparation for snATAC-seq

Single-cell libraries for snATAC-seq were generated using the GemCode Single-Cell Instruments and the Single Cell ATAC Library & Gel Bead Kit and ChIP Kit from 10X Genomics, following the manufacturer’s instructions. Samples were incubated at 37°C for 1 hour with 10 μl of transposition mix per reaction (7 μl of ATAC Buffer and 3 μl of ATAC Enzyme from 10X Genomics). Following the generation of nanoliter-scale GEMs, GEMs were reverse transcribed in a C1000 Touch Thermal Cycler (Bio-Rad) with the following program: 72°C for 5 minutes, 98°C for 30 seconds, 12 cycles of 98°C for 10 seconds, 59°C for 30 seconds, and 72°C for 1 minute, followed by holding at 15°C. After reverse transcription, single-cell droplets were broken, and the single-strand cDNA was isolated, cleaned up, and amplified. Amplified cDNA and final libraries were assessed using an Agilent BioAnalyzer with a High Sensitivity DNA Kit (Agilent Technologies). All libraries were sequenced on a NovaSeq 6000 platform (Illumina).

### Sample preparation for Bulk RNA-seq

Bulk RNA-Seq analysis was performed on distal ileums from control and NEC groups. Briefly, Total RNA was extracted using TRIZOL reagent (Invitrogen), and 4μg of total RNA were pooled and purified using the RNeasy Mini Kit (Qiagen) for each group. 10 μg of pooled total RNA were used to construct nondirectional RNA-Seq libraries (300 bp average insert size) using the TruSeq RNA-Seq Library Prep Kit v2 (Illumina, San Diego, CA) following the manufacturer’s instruction. Clusters were generated by using the cBot with libraries diluted to 20 pM and sequenced on a HiSeq2500 platform (Illumina). The sequencing was single end with 50 nt reads length.

### Sample Preparation and Analysis for Mass Spectrometry Proteomics

Tissues were lysed with Tissue Extraction Reagent I (ThermoFisher). Samples were homogenized at 30 Hz for 1.5 minutes and centrifuged at 13,000xg for 5 minutes. The supernatant was transferred to a new tube, and a BCA protein concentration assay was performed. A total of 25 µg of protein (in 5% SDS, 50 mM TEAB) was reduced with 20 mM DTT for 10 minutes at 95°C and alkylated with 40 mM iodoacetamide for 30 minutes in the dark. Samples were brought to a final concentration of 5% SDS, and phosphoric acid was added to reach a final concentration of 1.2%. Next, 165 µL of S-Trap protein binding buffer (90% methanol, 100 mM TEAB) was added to 27.5 µL of acidified lysate. The resulting mixture was passed through the micro column at 4000xg. The micro column was washed four times with the S-Trap protein binding buffer. Each sample was digested with 1 µg of trypsin (in 20 µL of 50 mM TEAB) for 1 hour at 47°C. Prior to elution, 40 µL of 50 mM TEAB (pH 8) was added to the column. Peptides were eluted by centrifugation at 4000xg. Elution was performed two more times using 40 µL of 0.2% formic acid and 40 µL of 50% acetonitrile + 0.2% formic acid. The eluted peptides were dried down and stored at −40°C.

For data-independent acquisition (DIA) LC-MS/MS, 250 ng of digested peptides were analyzed using nano-HPLC coupled to MS. A total of 250 ng of the sample was loaded onto Evotip Pure according to the manufacturer’s instructions. Peptides were eluted from the column (cat#: EV-1137, 15 cm x 150 µm with 1.5 µm beads) using the 30SPD pre-formed acetonitrile gradient generated by an Evosep One system and analyzed on a timsTOF Pro 2. The Evosep was coupled to the timsTOF Pro 2 using a 10 µm diameter emitter tip. The column toaster was set to 40°C. The total DIA protocol lasted 44 minutes. The MS1 scan had a mass range of 100–1700 Da in dia-PASEF mode. TIMS settings included an accumulation and ramp time of 100 ms, with a mobility range (1/K0) of 0.6 to 1.6 V·s/cm²and a cycle time of 2 seconds. For MS2, 1 mobility window with 2 ramps was used for 32 mass windows, each 29.8 Da wide with a 5 Da mass overlap. The mobility range was from 0.61/K0 to 1.451/K0, at a duty cycle of 100%, and a ramp rate of 9.52 Hz. 1+ ions were excluded from fragmentation using a polygonal filter. Auto calibration was turned off. Spectronaut v19.0 directDIA+ workflow was used to search the data with the Spectronaut-generated human spectral library (mouse_PDB_2023). Search parameters were set to default. Run-wise imputing was enabled, with global normalization applied.

### Data analysis for Bulk RNA-seq and Proteomics

For bulk RNA-seq data, raw RNA-seq count data were first transformed into counts per million (CPM). Genes with CPM above 1 expressed at least two samples were retained. Gene expression counts were normalized using the *voom* function from edgeR package. Differentially expressed genes (DEGs) were identified using the *eBayes* function from limma package. Genes with an adjusted p-value below 0.05 were considered significantly differentially expressed. For proteomics data, differentially expressed proteins (DEPs) were identified using the eBayes function from limma package. Proteins with an adjusted p-value below 0.05 were considered significantly differentially expressed.

### Data analysis for snRNA-seq

FASTQ sequence data were processed to obtain UMI counts in each cell using Cell Ranger (v7.2.0). A mouse reference genome (mm10) was used for gene alignment. snRNA-seq data analysis was performed using the Seurat package (v4.4.0) [78] in R. snRNA data was corrected for possible ambient RNA correction using DecontX [79]. Nuclei were discarded if they met any of the following criteria: (1) in the top 1% in terms of the number of genes, (2) with fewer than 200 genes or fewer than 500 UMIs, or (3) with more than 10% mitochondrial gene expression. Potential doublets were excluded using the scDblFinder package [80]. To select the collection of shared variable genes between samples, we first estimated the top 3,000 most variable genes per sample and then selected the top 3,000 most recurrent genes across all samples. PCA correction was performed using Harmony (v1.2.0) [21] with sample labels as covariates. UMAP was created with Seurat’s RunUMAP function using the first 30 principal components of Harmony’s PCA correction embedding. Differentially expressed genes (DEGs) were identified using Wilcoxon tests as implemented in Seurat’s FindAllMarkers function. Adjusted p-values were calculated using the Bonferroni correction for all features in the dataset. Genes with P_val_adj < 0.05 were considered DEGs. The final assignment of cells to major cell lineages was based on literature marker genes. To compare the cell count fold change in different conditions within each cluster, we performed the Monte Carlo permutation test (https://github.com/rpolicastro/scProportionTest) [81].

### Data analysis for snATAC-seq

The raw FASTQ data for each sample were processed using CellRanger for read filtering and alignment, barcode counting, identification of transposase cut sites, detection of accessible chromatin peaks, and cell calling, using the “cellranger-atac count” function (v2.1.0). The outputs from two runs of “cellranger-atac count” were further analyzed using “cellranger-atac aggr” to normalize input runs to the same median fragments per cell and to detect accessible chromatin peaks. For alignment, the mm10 reference genome was used.

The peaks called by cellranger-atac aggr were binarized and submitted to the R packages Signac (v1.1.0) [82] and Seurat for dimensional reduction and clustering analysis. Briefly, only cells meeting the following criteria were retained for further analysis: peak_region_fragments > 3000, peak_region_fragments < 300,000, pct_reads_in_peaks > 15, blacklist_ratio < 0.05, nucleosome_signal < 4, and TSS.enrichment > 2.5. Latent semantic indexing (LSI) was performed on the peaks of the remaining cells using the RunTFIDF and RunSVD functions of Signac to reduce dimensionality. To further remove batch effects, we performed integration using Harmony (v1.2.0) [21]. Major clusters were identified by using Seurat’s SNN graph clustering FindClusters at a resolution of 0.8. These same reduced dimensions were used as input into Seurat’s ‘RunUMAP’ with default parameters for visualization. To aid in cell type annotation in the scATAC-seq data, we applied Seurat’s integration framework to identify pairs of corresponding cells between the two modalities. This was achieved using the “FindTransferAnchors” and “TransferData” functions.

Cis analysis through peaks in the gene body and promoter was performed to generate gene activity scores, which were used to infer the cell types of clusters and integrate them with the snRNA-seq data. The gene body regions were extracted using the R package EnsDb.Mmusculus.v79, and the promoter region was defined as the 2000 base pairs upstream of the gene body. The gene activity score matrix was generated using the “GeneActivity” function of Signac, and only protein-coding genes were included.

The accessible chromatin peaks for each cell type were identified using the MACS2 method [22]. Differential chromatin accessibility analysis was performed using the “FindAllMarkers” function. Differentially accessible chromatin regions were identified with Bonferroni-adjusted p-values smaller than 0.05 and log2 fold-change values larger than 0.25. The genomic regions containing accessible chromatin peaks were annotated using ChIPSeeker (v1.38.0) [83] with the UCSC database on mm10.

Single-cell TF motif activity was estimated using the JASPAR 2020 database with the RunChromVAR wrapper in Signac. Differential TF activity between cell types was calculated using the “FindMarkers” function, with criteria set to log2 fold-change > 1 and Bonferroni-adjusted p-values < 0.05.

### Estimation of Cell Cycle Status and Module Score Definition

To assess the proliferation status of individual cells, we evaluated them using a characteristic gene set involved in the cell cycle, comprising 43 G1/S and 54 G2/M cell cycle genes, as previously described [84]. Cells manifesting high G1/S or G2/M scores were categorized as cycling, while those with low scores in both G1/S and G2/M were designated as non-cycling. A data-derived threshold, set at 2 Median Absolute Deviations (MADs) above the median, was applied to differentiate high from low scores.

For computing enterocyte differentiation programs along the villus axis, we used a gene signature [30] and employed the AddModuleScore() function from the Seurat R package to ascertain the extent to which individual cells expressed specific predefined expression programs. To compute gene set scores from the HARMARK gene set, signature scores were generated using the ScoreSignatures_UCell() function from the UCell package [85].

### Hotspot analysis

We determined transcriptional gene modules using the Hotspot package [41] in intestinal epithelial cells. We identified genes that were significantly autocorrelated with the PCA space using the “danb” observation model. A k-nearest neighbor graph was created using the create_knn_graph function with the parameters: n_neighbors = 30. We then computed pairwise local autocorrelations with Hotspot and clustered genes using these pairwise statistics with the create_modules function in Hotspot (minimum gene threshold of 20, FDR threshold of 0.05, core_only = False). This procedure identified 14 modules that were used for downstream analysis.

### CytoTRACE analysis

We performed CytoTRACE2 [31] analysis to infer the differentiation status of all epithelial clusters by using cytotrace2 function in R package CytoTRACE2. We applied MAGIC imputation [86] to normalized, log-transformed count matrices to denoise and recover missing transcript counts due to dropout. Function “magic” from R package Rmagic were used with parameters (t = 3, k = 5). Spearman’s correlation was computed between imputed gene expression and CytoTRACE2 scores to identify genes associated with stemness and differentiation.

### Celloracle analysis for epithelial cells

To construct gene regulatory networks (GRNs) and simulate the effects of transcription factor (TF) perturbation on epithelial cell gene expression, we utilized CellOracle (v0.18.0) [87] following the official documentation (https://morris-lab.github.io/CellOracle.documentation/). Initially, a base GRN was established using snATAC-seq data processed through Cicero [88] to identify accessible chromatin regions and potential TF binding sites. Subsequently, this network was pruned using snRNA-seq data and a Bagging Ridge model to refine regulatory interactions. Perturbation simulations involved setting the expression of specific TFs, such as Ezh2, to zero, enabling CellOracle to predict changes in gene expression and cellular transition trajectories. Pseudotime values, indicative of cellular differentiation states, were calculated from CytoTRACE2 scores using the formula: pseudotime = 1 - CytoTRACE2 scores, where higher values corresponded to more differentiated cells. This comprehensive methodology allowed us to accurately model and predict the impact of TF perturbations on gene expression dynamics within epithelial cells.

### Multi-modalities trajectories analysis

We employed the scMEGA workflow for integrating modalities and analyzing trajectories. snRNA-seq and snATAC-seq data were co-embedded using the “CoembedData” function. The PairCells function, employing the scOptmatch algorithm in “geodesic” pair mode, identified cell pairs with similar profiles across different modalities. Trajectory analysis was then conducted using the AddTrajectory function, with differentiation status ordered according to CytoTRACE2 scores, performed separately for absorptive and secretory lineages.

For variable peak identification along the trajectory, we first generated an accessibility matrix across all peaks using “GetTrajectory” function with log2Norm = TRUE. Peaks exhibiting variance greater than 0.9 were identified using “TrajectoryHeatmap” function, set with varCutOff = 0.9, returnMatrix = TRUE, and maxFeatures = 100000. Control and NEC group matrices were concatenated, selecting only peaks showing an absolute difference in magnitude of at least 0.2. TF motifs correlated with pseudotime were pinpointed using a correlation threshold of 0.5. Gene expression analysis involved setting log2Norm = TRUE in “GetTrajectory” function, filtering genes to include only those with an absolute magnitude difference of at least 0.5. We utilized the R package tradeSeq to identify genes differentially associated with pseudotime between the Control and NEC groups. Expression models were fitted using the “fitGAM” function, and genes linked to pseudotime were ascertained using the “startVsEndTest” function. Only genes with P-values less than 0.05 were retained for heatmap visualization.

### Enrichment analysis

GO enrichment analysis was conducted with the ‘enrichGO’ function (qvalueCutoff = 0.05) in the clusterProfiler v.4.10 package [89].

### Epithelial regulatory network inferred from VIPER algorithm

We employed a recently developed workflow to infer protein activity using the VIPER algorithm [42]. Briefly, we calculated protein activity profiles from snRNA-seq data for both the control and NEC groups separately. Metacells were aggregated from five cells per cell type. A regulatory network (interactome) was reverse-engineered from the resulting metacells using the ARACNe-AP algorithm with 200 bootstraps, a Mutual Information (MI) P-value threshold of P ≤ 10–8, and Data Processing Inequality (DPI) enabled. Regulatory proteins (RPs) were selected into manually curated protein sets to reconstruct the network (https://github.com/califano-lab/PISCES/). Regulons with fewer than 50 targets were excluded from the analysis, as recommended. The protein activity matrix was inferred using the VIPER algorithm, and downstream analyses, such as normalization and dimensionality reduction, were performed using the Seurat workflow.

### MERFISH library design and encoding probes

To deconvolute the spatial complexity of NEC pathogenesis, we utilized multiplexed error-robust fluorescence in situ hybridization (MERFISH) for targeted expression profiling in a spatial context. We designed a 500-gene panel targeting the most relevant pathways associated with NEC (Table S1), including those implicated in endothelial cells, immune cells, fibroblasts, smooth muscle cells, enteric neurons, and cells within the intestinal epithelium. The MERFISH library falls into the following categories: (1) cell-type markers and (2) genes known to regulate intestinal function. Established cell-type marker genes were used to distinguish different intestinal cell types and subtypes. The functional genes include those reported in NEC that regulate intestinal inflammation, vascularization, and regeneration. Each gene was assigned a unique 20-bit binary barcode which was used to decode sequential imaging data to assign detected probe barcode to the appropriate transcript.

### MERFISH tissue processing and imaging

MERFISH tissue processing and imaging were performed as previously reported [90]. All steps were performed under RNase-free conditions. Frozen embedded brains were cryo-sectioned at 10 μ m thickness at −21°C and mounted onto room temperature MERSCOPE beaded coverslips (Vizgen, Cat: 10500001). After adhering, sections were allowed to refreeze for 5-15 min, then fixed in 4% paraformaldehyde (PFA) diluted in 1X PBS for 15 min. Sections were washed three times with 1X PBS for 5 min each, then stored in 70% ethanol (EtOH) overnight at 4°C to permeabilize the tissue. Sections were stored in 70% EtOH for a maximum of 3 weeks. Sample preparation was performed using the sample preparation kit (Vizgen, Cat: 10400012) and Vizgen manufacturer instructions for unfixed tissue. First, sections were washed with 1X PBS followed by Sample Prep Wash Buffer (Vizgen, PN 20300001). Sections were incubated in Formamide Wash Buffer (Vizgen, PN 20300002) for 30 min at 37°C, then incubated in the gene panel mix for 42-46 hour (hr) at 37°C. Sections were then incubated two times in Formamide Wash Buffer for 30 min each at 47°C, and washed with sample prep wash buffer for at least 2 min. Sections were coated in gel embedding solution (0.05% w/v ammonium persulfate, 0.05% v/v N,N,N’,N’ - tetramethylethylenediamine in Gel Embedding Premix (Vizgen, PN 20300004)) and incubated at RT for 1.5 hr, cleared in 1:100 proteinase K in Clearing Premix (Vizgen, PN 20300003) at 37°C overnight or for a maximum of 7 days to clear lipids and proteins that may contribute to auto-fluorescence background noise. Prior to imaging, sections were washed two times with Sample Prep Wash Buffer, incubated in DAPI and PolyT Staining Reagent (Vizgen, PN 20300021) for 15 min on a rocking platform, incubated in Formamide Wash Buffer for 10 min, and washed again with Sample Prep Wash Buffer.

Imaging was performed on the MERSCOPE platform (Vizgen, Cat: 10000001) according to manufacturer instructions. Briefly, samples were loaded into the flow chamber of the instrument and the desired region for imaging was selected using a low-resolution mosaic of DAPI and PolyT stains. Samples were imaged at high-resolution with a 7-plane z-stack and 1.5 μ m spacing between adjacent z-planes to capture the entire 10 μ m thickness of the tissue sections. Samples were then automatically imaged according to MERSCOPE imaging presets.

### Preprocessing of MERFISH data

An RNA probe panel of 495 genes (Table S2) was selected to enable the identification of major cell types including growth factors, WNT signaling pathway genes, immune-related genes and cell type specific markers. After imaging, transcript barcodes were decoded and assigned to the appropriate gene. MERFISH images were segmented using Vizgen’s post-processing tool (VPT) and with a deep learning algorithm, CellPose 2.0 [91]. Cell boundaries were delineated based on pre-stained DAPI and PolyT signals. Individual RNA molecules were assigned to each cell based on their spatial location within the respective segmented cell boundaries. The decoded data were preprocessed by the following steps: (1) segmented ‘cells’ with a cell body volume less than 100 µm^3^ or larger than 4,000 were removed; (2) cells with total RNA counts of less than 10 or higher than 98% quantile, and cells with total RNA features less than 10, were removed; (3) to correct for the minor batch fluctuations in different MERFISH experiments, we normalized the total RNA counts per cell to a same value (500 in this case); (4) doublets were removed by Scrublet [92] and (5) the processed cell-by-gene matrix was transferred to gene-by-cell matrix and then loaded into Seurat V5 for downstream analysis [93]. The matrix was log-transformed by the Seurat standard pipeline. MERSCOPE Visualizer 2.3 (Vizgen) was used to manually annotate cells to their respective anatomical regions (crypt or villus) based on a mosaic pattern of DAPI and specific cell type markers. Segmented cells were overlapped onto histological staining, and cells were manually selected to their respective anatomical region based on histological definitions of crypt and villus. A minimum of two layers of cells from the epithelial barrier were selected to encompass the neighbourhood of cells residing in the respective anatomical regions.

To transfer labels from snRNA-seq data to MERFISH data, we employed Seurat’s label transfer framework. We used Seurat’s *FindTransferAnchors* function and *TransferData* function to transfer the snRNA-seq cell type labels onto the MERFISH dataset. To impute gene expression in MERFISH data, we utilized the iSpatial algorithm [94], a method specifically designed for spatial transcriptomics data by leveraging gene expression profiles information from the snRNA-seq data. To perform differential expression analysis using the raw MERFISH count data, we applied Seurat’s *FindMarkers* function. Genes with adjusted p-value < 0.05 were considered as significantly differentially expressed genes (DEGs). GO enrichment analysis was then conducted by using the clusterProfiler package [89].

### Statistics

All analyses were performed with GraphPad prism 9. All results were performed at least in triplicate and expressed as mean ±SD. P-values were determined using nonparametric unpaired Student’s t-tests (Mann–Whitney U test) or using one-way ANOVA with post hoc Tukey’s test as appropriate. P <0.05 was considered as statistically significant.

## Data availability

All relevant data supporting the key findings of this study, as well as the raw image data generated in this study, are available within the paper and its Supplementary Information or from the corresponding author upon reasonable request. The raw sequence data reported in this paper have been deposited in the Gene Expression Omnibus (GEO) that are publicly accessible at GSE278973.

## Author Contributions

YX, AZ, HZ, and BL were involved in the study design; AZ, GB, FB, MY, CL, DL, CW, FM, SW, and YT were involved in the animal data collection, analysis, and interpretation; HL and HZ were involved in sample preparation for RNA sequencing analysis; YX, JY, JH, and BL were involved in single-cell RNA sequencing data analysis, and interpretation; NT and BK were involved in MERFISH data collection and interpretation, AM and PDO were involved in Ezh2 experiments; PDC, AP, HZ, and BL provided advice and supervision. All the authors reviewed and approved the final manuscript. YX, AZ, and HL contributed equally as the first author of this work. AP, HZ, and BL contributed equally as the corresponding authors.

## Acknowledgements

The authors wish to thank Laura McGary and Cassandra Wong of the Network Biology Collaborative Centre Proteomics Facility (RRID: SCR_025375) at the Lunenfeld-Tanenbaum Research Institute for global proteome analysis. The facility is supported by the Canada Foundation for Innovation and the Ontario Government.

