## Supplementary Figures and Figure Legend for "Molecular and Epigenetic Pathways Underlying Epithelial Damage and Repair in Necrotizing Enterocolitis via Multi-omics Approach"

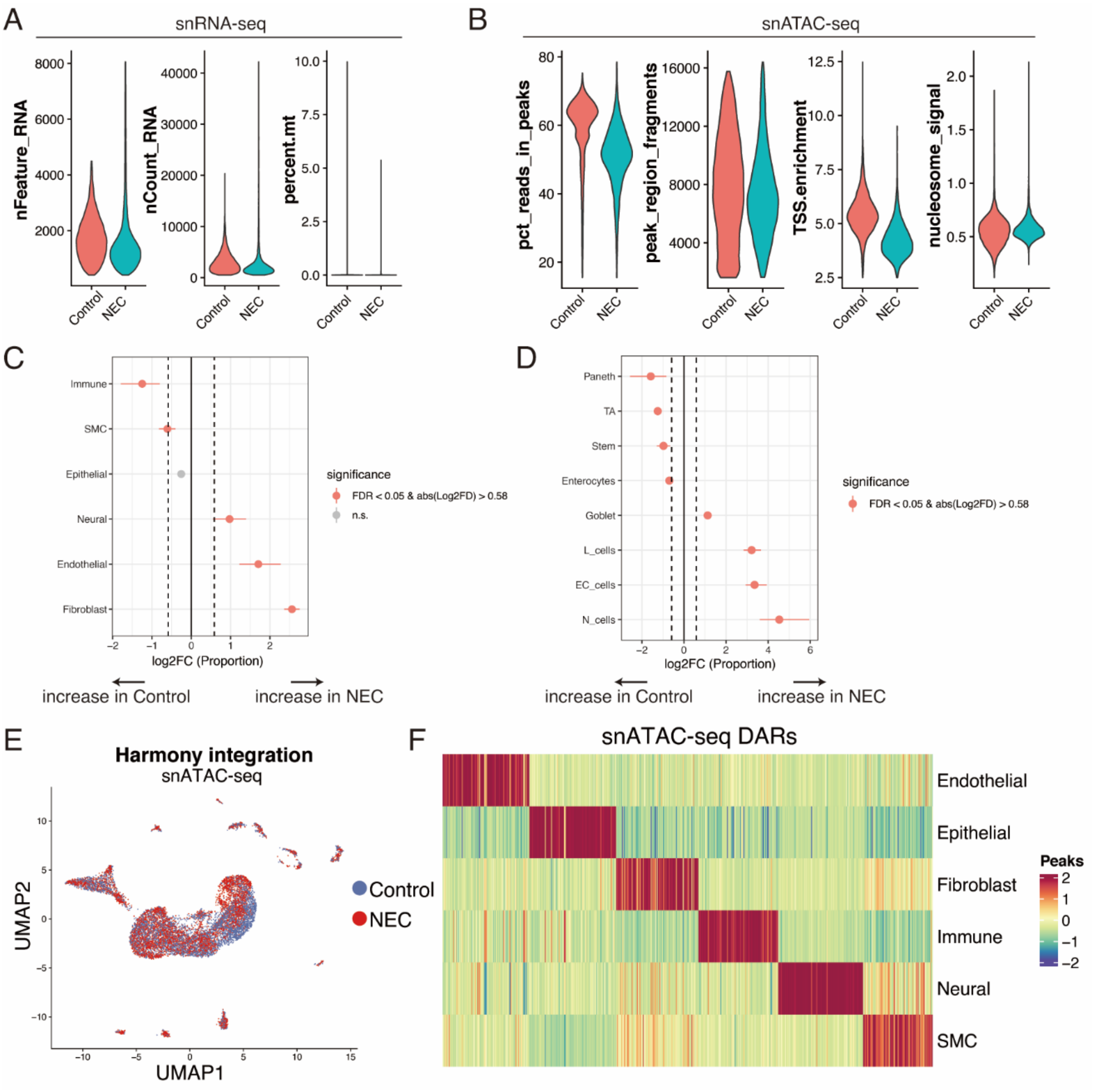


**Figure S1. Quality control of snRNA-seq, snATAC-seq data.**

(A) Violin plots showing quality control metrics of snRNA-seq data for high-quality cells. (B) Violin plots showing quality control metrics of snATAC-seq data for high-quality cells. (C) UMAP view of snATAC-seq data after harmony integration. (D) Differentially accessible regions (DARs) identified per cell type. (E) Cell proportion fold change in NEC vs control for major cell compartments. (F) Cell proportion fold change in NEC vs control for epithelial cell types.

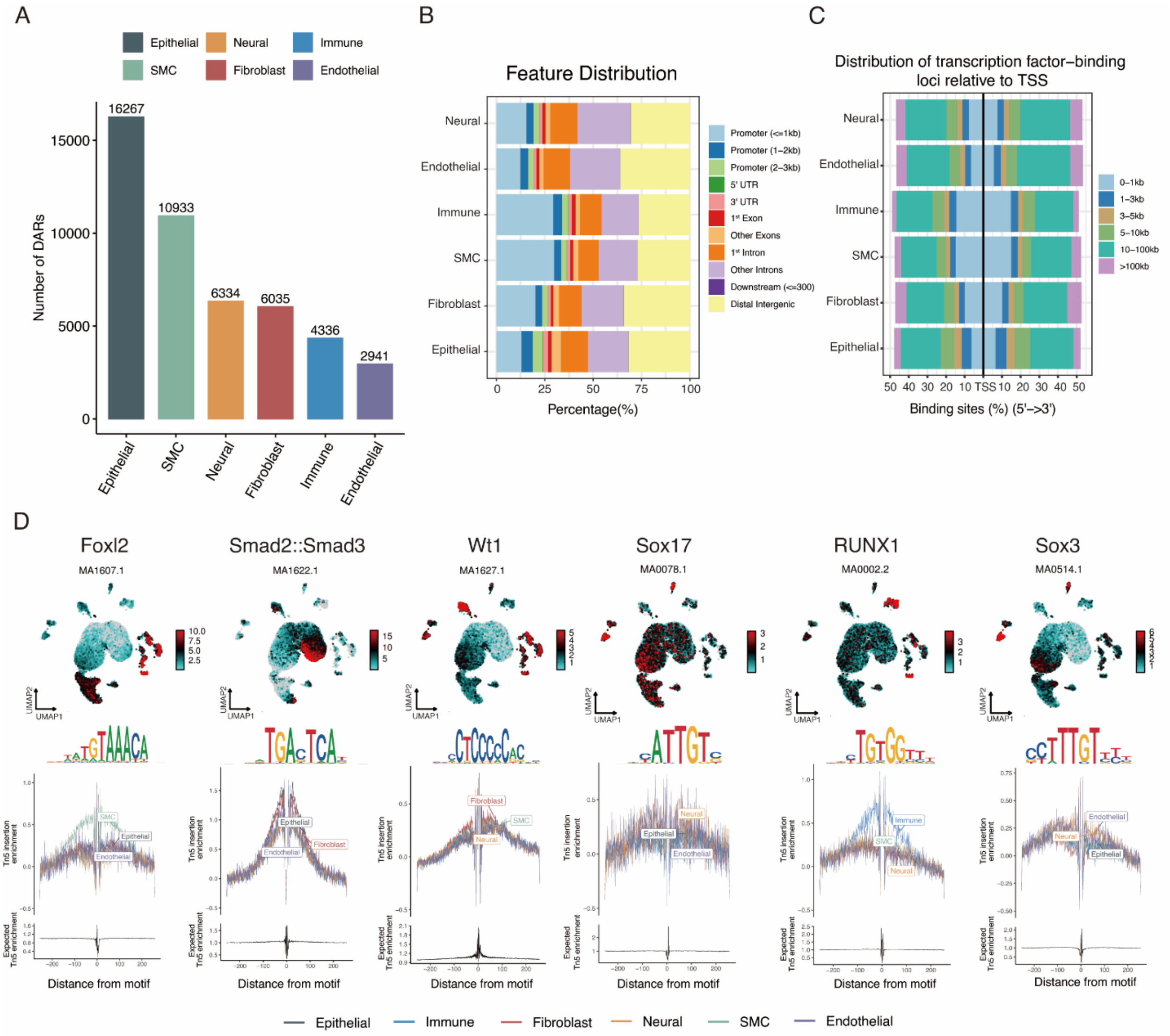


**Figure S2. Characterization of DARs in all compartments from snATAC-seq data.**

(A) Bar graphs showing the number of differentially accessible regions (DARs) per cell type. (B) Bar graphs showing the annotated genomic locations for each cell type. (C) Bar graphs showing the distance of the transcription start site of DARs for each cell type. (D) TF motif analysis showing enriched TF motifs in each cell type.

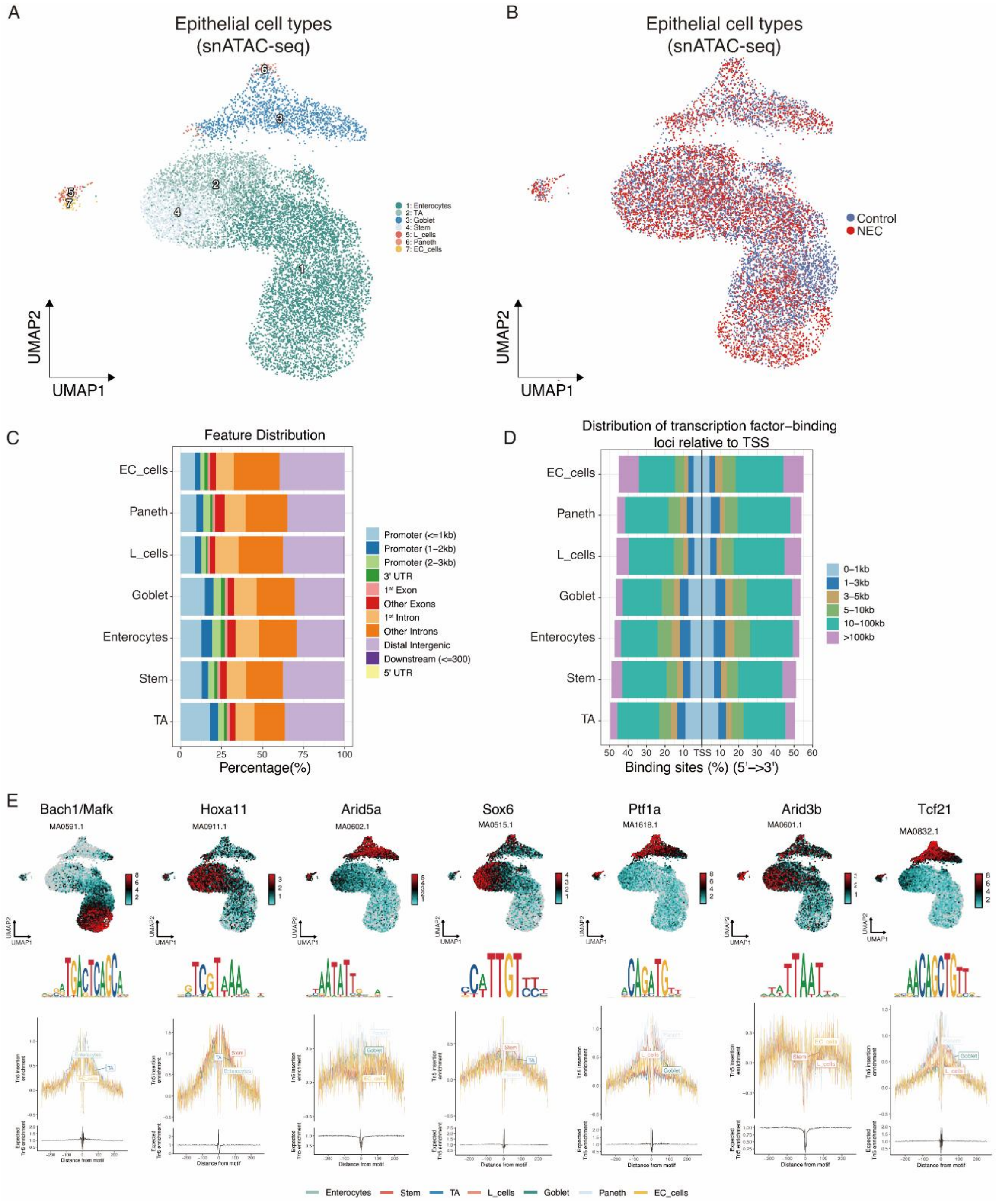


**Figure S3. Characterization of DARs in the epithelial compartment from snATAC-seq data.**

(A) UMAP view of epithelial cell types in snATAC-seq data. (B) UMAP view of sample group in snATAC-seq data. (C) Bar graphs showing the annotated genomic locations for each epithelial cell type. (D) Bar graphs showing the distance of the transcription start site of DARs for each cell type in epithelial cells. (E) TF motif analysis showing enriched TF motifs in each epithelial cell type.

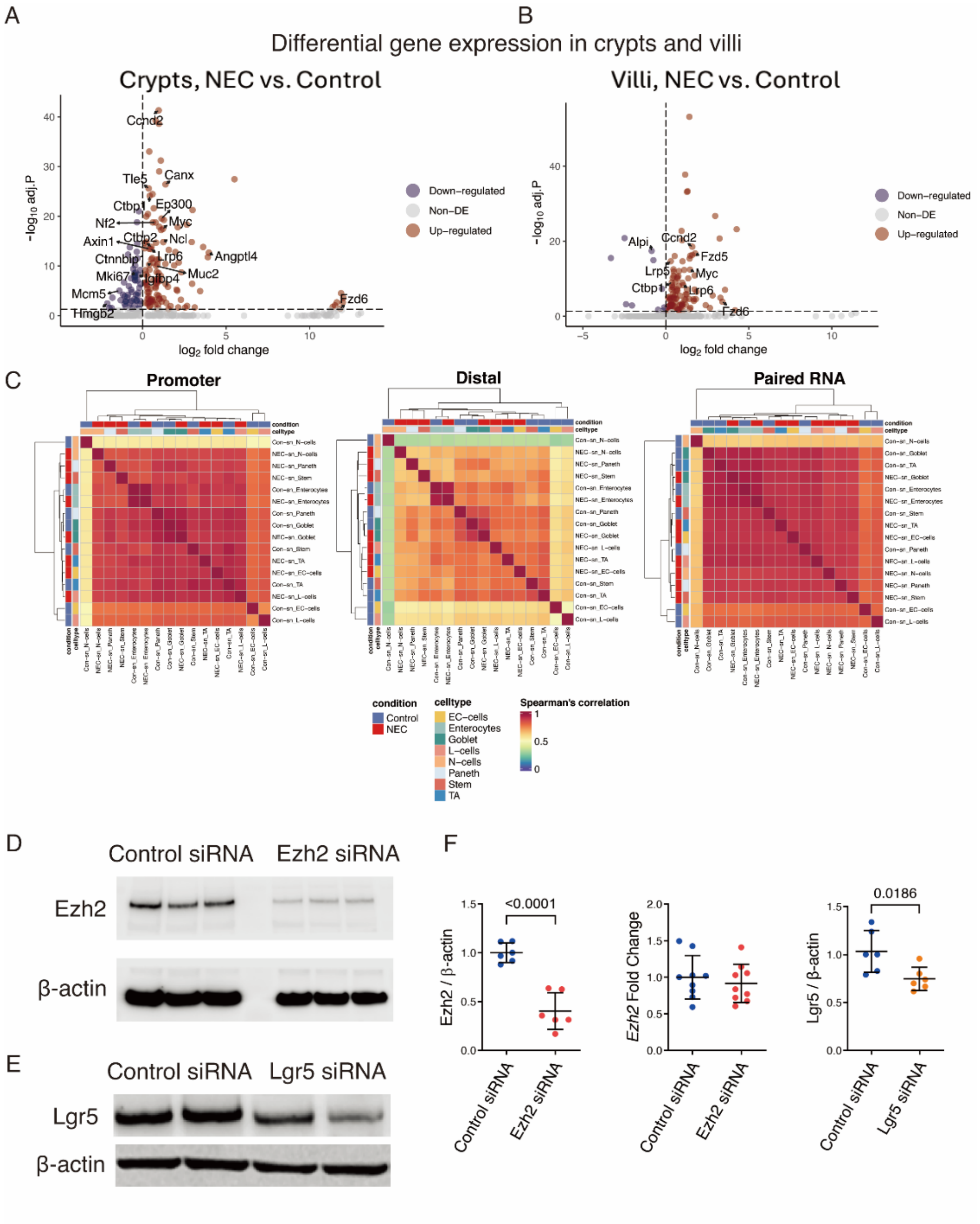


**Figure S4. Comprehensive analysis of gene expression, chromatin accessibility, and siRNA transfection in intestinal epithelial cells**

(A) Volcano plots showing differentially expressed genes that are upregulated and downregulated in NEC within the crypt and (B) villus regions based on MERFISH datasets. (C) Pairwise Pearson correlation of aggregate single-cell chromatin accessibility profiles associated with gene promoters, area distal from the promoters, and paired gene expression, aggregated by cell type and condition. (D) Representative micrographs of EZH2 gel blot in IEC-18 cells treated with Ezh2 siRNA compared to cells treated with a negative control vector, with β-actin as loading control. (E) Representative micrographs of LGR5 gel blot in IEC-18 cells treated with Lgr5 siRNA compared to cells treated with a negative control vector, with β-actin as loading control. (F) Qualification of

EZH2 protein expression and gene expression in IEC-18 cells treated with Ezh2 siRNA compared to cells treated with a negative control vector and qualification of LGR5 protein expression in IEC-18 cells treated with Lgr5 siRNA compared to cells treated with a negative control vector. Experiments were repeated independently 3 times, yielding similar results. Each dot represents the average value for each individual. Data are presented as mean ± SD and compared using one-way ANOVA with post-hoc tests. The primary p-value is indicated directly on the graph.
