## Supplementary Tables for "Molecular and Epigenetic Pathways Underlying Epithelial Damage and Repair in Necrotizing Enterocolitis via Multi-omics Approach"

**Table S1. Primer sequences used for quantitative PCR.**

| Gene | Forward Sequence | Reverse Sequence |
|---|---|---|
| *TNF-α (m)* | *TTCCGAATTCACTGGAGCCTCGAA* | *TGCACCTCAGGGAAGAATCTGGAA* |
| *IL-6 (m)* | *TGGCTAAGGACCAAGACCATC* | *TTCTGACCACAGTGAGGAATGTC* |
| *LGR5 (m)* | *CGAGCCTTACAGAGCCTGATACC* | *TTGCCGTCGTCTTTATTCCATTGG* |
| *GAPDH (m)* | *TGAAGCAGGCATCTGAGGG* | *CGAAGGTGGAAGAGTGGGAG* |
| *SOX9 (m)* | *CTGGAGGCTGCTGAACGAGAG* | *CGGCGGACCCTGAGATTGC* |
| *PCNA(m)* | *TTTGAGGCACGCCTGATCC* | *GGAGACGTGAGACGAGTCCAT* |
| *EZH2 (m)* | *TCAAAACCGCTTTCCTGG* | *TGTCCCAATGGTCAGCA* |
| *KLF4 (m)* | *ACCTGGCGAGTCTGACATGG* | *TCCTCACGCCAACGGTTAGT* |
| *KLF5 (m)* | *ACTGCCCTCGGAGGAGCTGG* | *ATGCTCTGAAATTATCGGAACTG* |
| *MUC2 (m)* | *GAACGGGGCCATGGTCAGCA* | *CATAATTGGTCTGCATGCC* |
| *LYZ1(m)* | *GAGAACCGAAGCACCGACTATG* | *CGGTTTTGACATTGTGTTCGC* |

**Table S2. Gene panel for MERFISH data**

| Genes | | | | | | | | | |
|---|---|---|---|---|---|---|---|---|---|
| *Acta1* | *Canx* | *Csf1* | *Fbxw2* | *Hmgb1* | *Kdm6b* | *Nhp2* | *Pparg* | *Slc17a7* | *Tnf* |
| *Acvr2a* | *Car4* | *Csf1r* | *Fbxw4* | *Hmgb2* | *Kdr* | *Nkd1* | *Ppp2ca* | *Slc1a3* | *Tnfrsf19* |
| *Acvrl1* | *Car7* | *Csf2* | *Fcgr2b* | *Hmgcs2* | *Klf4* | *Nkx2.3* | *Ppp2r1a* | *Slc26a3* | *Tnfrsf1a* |
| *Adamts3* | *Casp1* | *Csf3* | *Ffar4* | *Hopx* | *Kmt2a* | *Nlgn2* | *Prdx1* | *Smoc2* | *Tnfrsf1b* |
| *Adcy8* | *Casp8* | *Csnk1a1* | *Fgf21* | *Hrg* | *Kremen1* | *Nlk* | *Prf1* | *Socs1* | *Tnfsf12* |
| *Adgra2* | *Ccbe1* | *Csnk1d* | *Fgf4* | *Hsf1* | *Krt18* | *Nme1* | *Prkaa1* | *Socs3* | *Tnfsf13* |
| *Adipoq* | *Ccl22* | *Csnk1g1* | *Fkbp1a* | *Hspb1* | *Krt7* | *Nos2* | *Prkca* | *Sox11* | *Tnni3* |
| *Adora1* | *Ccl3* | *Csnk2a1* | *Fkbp5* | *Htatip2* | *L3mbtl3* | *Notch1* | *Prkd1* | *Sox17* | *Top2a* |
| *Adora2a* | *Ccn4* | *Ctbp1* | *Flt4* | *Hvcn1* | *Lcn2* | *Notch4* | *Prkd2* | *Sox2* | *Trem1* |
| *Ager* | *Ccna2* | *Ctbp2* | *Fn1* | *Id2* | *Ldha* | *Nox1* | *Prok2* | *Sox3* | *Trem2* |
| *Aggf1* | *Ccnd1* | *Ctnnb1* | *Fosl1* | *Ifnb1* | *Lef1* | *Nox4* | *Prom1* | *Sox9* | *Trib1* |
| *Ahnak* | *Ccnd2* | *Ctnnbip1* | *Foxc1* | *Ifng* | *Lgals3* | *Nppb* | *Prox1* | *Sp8* | *Trp53* |
| *Aif1* | *Ccnd3* | *Ctsb* | *Foxg1* | *Ifngr1* | *Lgr5* | *Npr1* | *Psmb8* | *Sparc* | *Trp73* |
| *Aifm1* | *Ccr2* | *Ctsl* | *Foxj1* | *Ifngr2* | *Lif* | *Npr2* | *Ptgs2* | *Spdef* | *Tspan12* |
| *Alcam* | *Ccr3* | *Cx3cr1* | *Foxn1* | *Igfbp4* | *Lmna* | *Nr3c1* | *Ptk2b* | *Sphk1* | *Tspan2* |
| *Alpi* | *Ccr5* | *Cxadr* | *Foxo3* | *Iigp1* | *Lrig1* | *Nrp1* | *Ptp4a3* | *Sphk2* | *Tspo* |
| *Amigo2* | *Cd109* | *Cxcl10* | *Foxo4* | *Il10* | *Lrp5* | *Nrp2* | *Ptpn2* | *Spi1* | *Tubb3* |
| *Amot* | *Cd14* | *Cxcl16* | *Frat1* | *Il10ra* | *Lrp6* | *Nrros* | *Ptx3* | *Spib* | *Tubb4b* |
| *Ang* | *Cd163* | *Cxcr5* | *Frzb* | *Il10rb* | *Lum* | *Nus1* | *Pygo1* | *Spink5* | *Tubb5* |
| *Angptl3* | *Cd24a* | *Cxxc4* | *Fshb* | *Il11* | *Ly6a* | *Olfm4* | *Pyy* | *Spink6* | *Txn1* |
| *Angptl4* | *Cd4* | *Cybb* | *Fzd1* | *Il12a* | *Ly6c1* | *Olig2* | *Rb1* | *Sprr1a* | *Vcam1* |
| *Anxa1* | *Cd44* | *Cyc1* | *Fzd2* | *Il12b* | *Ly6d* | *Osmr* | *Rest* | *Spry2* | *Vegfa* |
| *Anxa3* | *Cd63* | *Cysltr2* | *Fzd3* | *Il13* | *Lyve1* | *P2rx7* | *Retnla* | *Srebf1* | *Vegfb* |
| *Apc* | *Cd68* | *Daam1* | *Fzd4* | *Il13ra1* | *Lyz2* | *P2ry12* | *Rhob* | *Srgn* | *Vegfc* |
| *App* | *Cdc42* | *Dab2ip* | *Fzd5* | *Il15* | *Mbp* | *P2ry2* | *Rhou* | *Stab1* | *Vegfd* |
| *Areg* | *Cdh1* | *Dclk1* | *Fzd6* | *Il17f* | *Mcm5* | *P2ry6* | *Ripk1* | *Stab2* | *Vim* |
| *Arg1* | *Cdh13* | *Dcn* | *Fzd7* | *Il18* | *Mecp2* | *Parp1* | *Rnh1* | *Steap4* | *Wif1* |
| *Ascl1* | *Cdh15* | *Dcx* | *Fzd8* | *Il1a* | *Mif* | *Pax6* | *Robo1* | *Stk11* | *Wnt1* |
| *Ascl2* | *Cdh17* | *Defb1* | *Gab1* | *Il1b* | *Mki67* | *Pck1* | *Robo4* | *Tcf4* | *Wnt10a* |
| *Aspg* | *Cdhr5* | *Dixdc1* | *Gab3* | *Il1r1* | *Mmp7* | *Pcna* | *Ror2* | *Tcf7* | *Wnt11* |
| *Ass1* | *Cdk6* | *Dkk1* | *Gad2* | *Il1r2* | *Mmp9* | *Pdgfa* | *Rps6kb1* | *Tcf7l1* | *Wnt16* |
| *Atoh1* | *Cdkn1a* | *Dll1* | *Gata2* | *Il1rn* | *Mptx2* | *Pdgfb* | *Runx1* | *Tert* | *Wnt2* |
| *Atp11b* | *Cdo1* | *Dnmt1* | *Gba* | *Il2* | *Mrc1* | *Pdgfra* | *Runx2* | *Tet1* | *Wnt2b* |
| *Atpif1* | *Cebpa* | *Dnmt3a* | *Gbp2* | *Il3* | *Msi1* | *Pdgfrb* | *S100a10* | *Tgfb1* | *Wnt3* |
| *Axin1* | *Cebpe* | *Dnmt3b* | *Gcgr* | *Il31ra* | *Msn* | *Peg3* | *S1pr3* | *Tgfb2* | *Wnt3a* |
| *Axl* | *Chga* | *Dvl1* | *Gfap* | *Il34* | *Msr1* | *Pf4* | *Scg2* | *Tgfb3* | *Wnt4* |
| *B3gnt5* | *Chgb* | *Dvl2* | *Ggta1* | *Il4* | *Muc1* | *Pfkfb1* | *Scg3* | *Tgfbi* | *Wnt5a* |
| *Bcl11b* | *Chil3* | *Egf* | *Gjb2* | *Il5* | *Muc2* | *Pgf* | *Sell* | *Tgfbr1* | *Wnt5b* |
| *Bcl9* | *Chrna7* | *Eif2ak1* | *Glp1r* | *Il6* | *Muc3* | *Pglyrp1* | *Sema6a* | *Tgfbr2* | *Wnt6* |
| *Bmi1* | *Clca1* | *Emcn* | *Glp2r* | *Il7* | *Muc4* | *Phlda1* | *Senp2* | *Timp1* | *Wnt7a* |
| *Bmp4* | *Clcf1* | *Emp1* | *Grip1* | *Inha* | *Myc* | *Pik3ca* | *Serpina3n* | *Timp2* | *Wnt7b* |
| *Btg1* | *Cldn3* | *Eno1* | *Gsk3a* | *Inhba* | *Myh10* | *Pik3cb* | *Serpinb1a* | *Tkt* | *Wnt8a* |
| *Btg2* | *Cldn4* | *Ep300* | *Gsk3b* | *Itgam* | *Myh9* | *Pik3cd* | *Serpine1* | *Tle1* | *Wnt9a* |
| *Btrc* | *Clec7a* | *Epgn* | *Gsn* | *Itgb1* | *Myo1c* | *Pitx2* | *Serpinf2* | *Tle2* | *Xdh* |
| *C1galt1* | *Clu* | *Ephb2* | *Gsta4* | *Itgb2* | *Ncf1* | *Pkm* | *Sfrp1* | *Tle5* | *Zbtb46* |
| *C1qc* | *Col4a2* | *Erap1* | *Havcr2* | *Itgb4* | *Ncl* | *Pla2g2a* | *Sfrp4* | *Tlr2* | |
| *C1ra* | *Col4a3* | *Ezh2* | *Hcls1* | *Jcad* | *Neurod1* | *Plg* | *Shh* | *Tlr4* | |
| *C3* | *Cps1* | *Fabp5* | *Hdac1* | *Jun* | *Nf1* | *Pml* | *Sirpa* | *Tm4sf1* | |
| *C5ar1* | *Creb1* | *Fadd* | *Hdac2* | *Kat6b* | *Nf2* | *Pnliprp2* | *Sirt1* | *Tmem119* | |
| *Cadm4* | *Cry1* | *Fbxw11* | *Hes6* | *Kdm1a* | *Nfkb1* | *Porcn* | *Slc10a6* | *Tmem37* | |
